# Structures of the PKA RIα holoenzyme with the FLHCC driver J-PKAcα or wild type PKAcα

**DOI:** 10.1101/505347

**Authors:** Baohua Cao, Tsan-Wen Lu, Juliana A. Martinez Fiesco, Michael Tomasini, Lixin Fan, Sanford M. Simon, Susan S. Taylor, Ping Zhang

## Abstract

Fibrolamellar hepatocellular carcinoma (FLHCC) is driven by J-PKAcα, a kinase fusion chimera of the J-domain of DnaJB1 with PKAcα, the catalytic subunit of Protein Kinase A (PKA). Here we report the crystal structures of the chimeric fusion RIα_2_:J-PKAcα_2_ holoenzyme formed by J-PKAcα and the PKA regulatory (R) subunit RIα, and the wild type (wt) RIα_2_:PKAcα_2_ holoenzyme. The chimeric and wt RIα holoenzymes have quaternary structures different from the previously solved wt RIβ and RIIβ holoenzymes. The chimeric holoenzyme shows an isoform-specific interface dominated by antiparallel interactions between the N3A-N3A’ motifs of RIα that serves as an anchor for RIα structural rearrangements during cAMP activation. The wt RIα holoenzyme showed the same configuration as well as a distinct second conformation. In the structure of the chimeric fusion RIα_2_:J-PKAcα_2_ holoenzyme, the presence of the J-domain does not prevent formation of the holoenzymes, and is positioned away from the symmetrical interface between the two RIα:J-PKAcα heterodimers in the holoenzyme. The J-domains have significantly higher temperature factors than the rest of the holoenzyme, implying a large degree of conformational flexibility. Furthermore molecular dynamics simulations were applied to analyze the conformational states of chimeric fusion and wt RIα holoenzymes, and showed an ensemble of conformations in the majority of which the J-domain was dynamic and rotated away from the R:J-PKAcα interface. Thus, rather than affecting the interactions with the regulatory subunits, the fusion of the J-domain to the PKAcα alters the conformational landscape of the chimeric fusion holoenzymes and potentially, as result, the interactions with other molecules. The structural and dynamic features of these holoenzymes enhance our understanding of the fusion chimera protein J-PKAcα that drives FLHCC as well as the isoform specificity of PKA.

## Introduction

FLHCC is a rare liver cancer that predominantly affects adolescent and young adults with no history of liver disease (Craig et al., 1980; Eggert et al., 2013; Honeyman et al., 2014; Kakar et al., 2005; Lalazar and Simon, 2018; Torbenson, 2012). It does not respond well to chemotherapy and the overall five year survival rate of FLHCC patients is only 30-45% (El-Serag and Davila, 2004; Kakar et al., 2005; Katzenstein et al., 2003; Lim et al., 2014; Mavros et al., 2012; Weeda et al., 2013). The chimeric gene *DNAJB1-PRKACA*, ubiquitously and exclusively found in almost all FLHCC patients, is the result of a ~400 kb deletion in one copy of chromosome 19 (Darcy et al., 2015; Engelholm et al., 2017; Honeyman et al., 2014; Kastenhuber et al., 2017; Oikawa et al., 2015; Riggle et al., 2016a, 2016b; Simon et al., 2015). This produces an enzymatically active chimeric protein J-PKAcα. The tumor is driven not by the deletion but by the formation of the J-PKAcα fusion protein, and the tumorigenicity of J-PKAcα is dependent on its kinase activity (Kastenhuber et al., 2017). The fusion chimera protein has the first 69 residues of the N-terminus of DnaJB1, namely the J-domain, and the C-terminal 336 residues of PKAcα (Cheung et al., 2015; Honeyman et al., 2014) (Figure 1A). In its inactive state in cells, PKA exists as a holoenzyme composed of two catalytic subunits and one regulatory (R) subunit homodimer (Taylor et al., 2012). Cyclic adenosine monophosphate (cAMP) binding to the R subunits unleashes the PKAcα activity. Each R subunit is composed of an N-terminal dimerization/docking (D/D) domain followed by a flexible linker and two tandem highly conserved cyclic nucleotide-binding domains (CNB-A and CNB-B) (Figure 1A). There are four functionally non-redundant R isoforms, RIα, RIβ, RIIα, and RIIβ with similar domain organization (Taylor et al., 2012). The engineered R:PKAcα heterodimers where one PKAcα subunit is bound to a truncated monomeric form of the R subunit all appear to be very similar (Boettcher et al., 2011; Ilouz et al., 2012; Zhang et al., 2012). However, when the two R:PKAcα heterodimers, linked to the D/D domain by the flexible linkers are assembled into holoenzymes, each forms a unique symmetry-related interface between the two heterodimers and thus creates isoform-specific quaternary structures, as shown by the solved structures of the RIβ and RIIβ holoenzymes and the RIα holoenzyme model (Boettcher et al., 2011; Ilouz et al., 2012; Zhang et al., 2012) (Figure S1). Among the four R isoforms, RIα can be considered as a master regulator for PKA signaling in mammalian cells. Deletion of RIα, for example, is embryonically lethal in mice and leads to unregulated PKA activity (Amieux et al., 1997). RIα also compensates when other R subunits are depleted or when PKAcα is overexpressed (Amieux and McKnight, 2002). It is the only upregulated R isoform in FLHCC cancer cells (Riggle et al., 2016b; Simon et al., 2015). Haploinsufficiency of RIα leads to a wide range of disease states, including Carney Complex (CNC) disease (Linglart et al., 2012; Park et al., 2012; Veugelers et al., 2004) as the other R subunit isoforms cannot compensate(Greene et al., 2008). Interestingly, recent studies (Graham et al., 2017; Terracciano et al., 2004) identified three individual patients with FLHCC and a personal history of CNC disease although the majority of CNC patients have no history of FHLCC.

**Figure 1.**
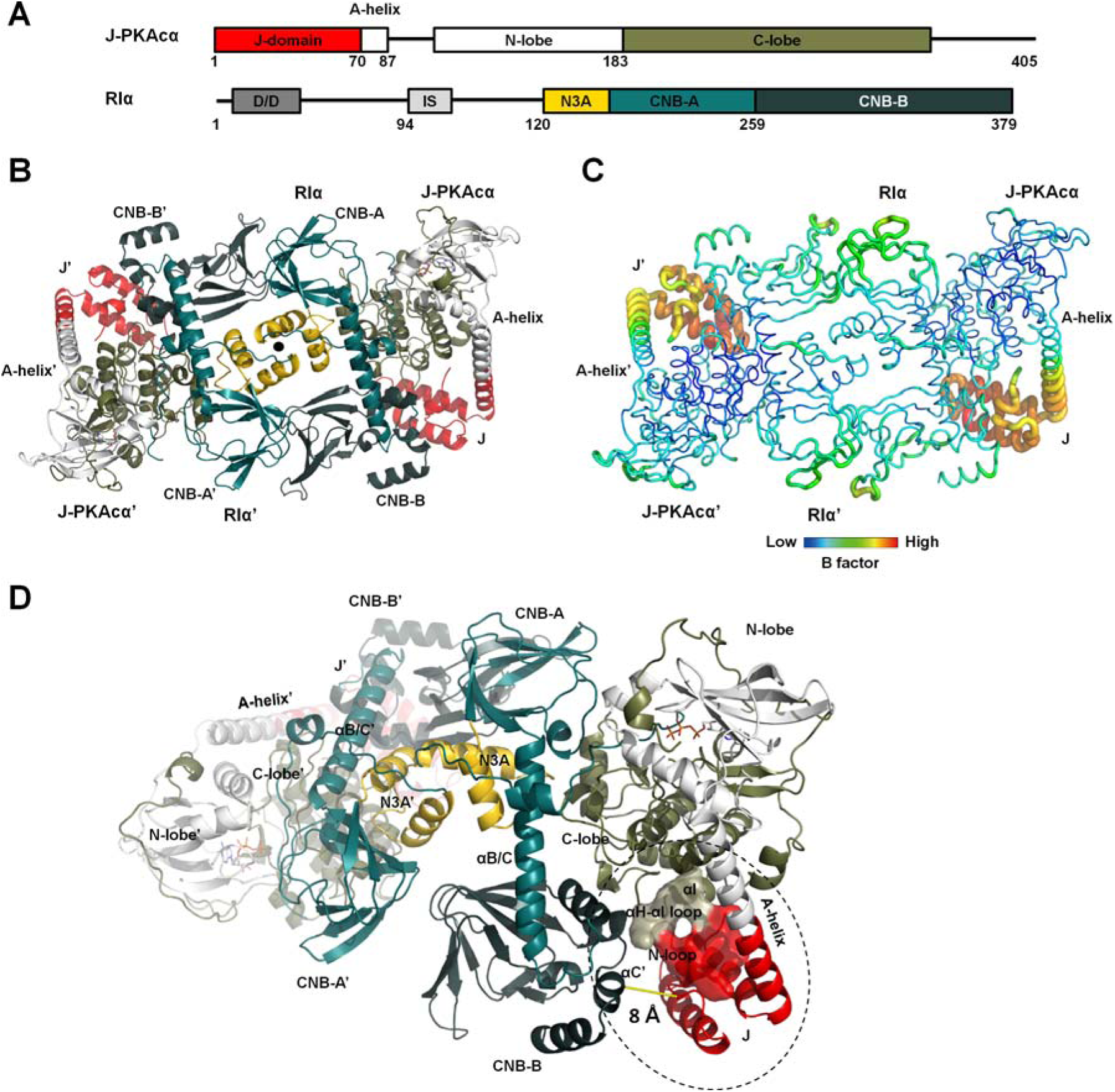
Overall structure of the chimeric RIα_2_:J-PKAcα_2_ holoenzyme. (A) Domain organization and color coding of J-PKAcα and RIα subunits. (B) Structure of the holoenzyme. One heterodimer is labeled as RIα:J-PKAcα and its two-fold symmetry mate is labeled as RIα’:J-PKAcα’. The two-fold axis position is shown as a solid black circle. (C) B factor analysis of the holoenzyme. (D) The J-domain is in close proximity to the CNB-B domain of RIα.

Structural studies of the J-PKAcα chimera (Cheung et al., 2015) showed that it has all the structural hallmarks of wt PKAcα with the conserved bilobal kinase core shared by all kinase superfamily members. The only structural alteration is the fused J-domain, which replaces the myristylation motif (residues 1-14). In the crystal structure the J-domain is tucked underneath the C-lobe of the conserved kinase core (Cheung et al., 2015). However, in molecular dynamics (MD) simulations and NMR assays the fused J-domain explores a large diffusional space (Tomasini et al., 2018). Though J-PKAcα is overexpressed relative to PKAcα in FLHCC cells (Honeyman et al., 2014; Simon et al., 2015), overexpression of PKAcα alone is insufficient to recapitulate the oncogenic effect of J-PKAcα (Kastenhuber et al., 2017). Compensatory expression of RIα mRNA and protein were detected in FLHCC tumors while both the mRNA and protein levels of RIIβ are down-regulated (Riggle et al., 2016b; Simon et al., 2015). J-PKAcα can interact with truncated RIα and RIIβ to form R:J-PKAcα heterodimers *in vitro* (Cheung et al., 2015), suggesting that both wt PKAcα and the chimeric J-PKAcα can form holoenzymes. To understand how PKA signaling might be disrupted by the FLHCC chimera it is essential to appreciate the architecture of the chimeric and wt holoenzymes as well as knowledge of their dynamics. In this study, we show that the J-PKAcα chimera is inhibited by full-length RIα and capable of forming the canonical holoenzyme with activation still under the control of cAMP. We report the crystal structures of the oncogenic RIα chimeric holoenzyme and the wt holoenzyme at 3.66 Å and 4.75 Å resolution, respectively (Figure S2). To explore whether the addition of the J-domain affects the conformational landscape of each holoenzyme, we furthermore report on MD simulations of the chimeric and wt RIα holoenzymes. We found states where the J-domain of J-PKAcα is able to interact with the C-terminal CNB-B domain of the RIα subunits; however, in the majority of MD states, the J-domain was dynamic and rotated away from the R:PKAcα interface. Altogether, these structural and dynamic descriptions of the driver of FLHCC enhance our understanding the molecular mechanism of this disease as well as our understanding of the dynamic allosteric mechanisms that couple cAMP binding to PKA activation.

## Results

### Overall structure of the FLHCC driver RIα_2_:J-PKAcα_2_ chimeric fusion holoenzyme

The complex of the full-length RIα and J-PKAcα chimera was formed *in vitro* by mixing the individually purified subunits followed by gel filtration (Figure S3). The full-length holoenzyme structure was determined at 3.66 Å resolution (Figures. 1B, S4 and Table 1). Each asymmetric unit (ASU) contains one holoenzyme molecule consisting of an RIα homodimer and two chimeric J-PKAcα subunits, thus the chimeric holoenzyme has the same stoichiometry as the previously published wt holoenzymes (Taylor et al., 2012). The presence of the J-domain does not prevent formation of the holoenzymes, and is positioned away from the symmetrical interface between the two RIα:J-PKAcα heterodimers in the holoenzyme. The J-domain can be easily accommodated spatially in the holoenzyme complex; there appears to be no steric constraints. The interface between the two heterodimers in the chimeric holoenzyme is strictly two-fold symmetry-related and created solely by the two RIα subunits, which pack against each other in an antiparallel orientation that includes a four-helical bundle involving the N3A motifs of the RIα subunits (Figure 1B). The PKAcα part of the chimera is almost identical to the PKI-bound wt PKAcα structure (Zheng et al., 1993), with a Cα root mean square deviation (RMSD) of 0.42 Å. The only structural alteration is a more linear and extended A-helix fused with the J-domain (Figure S5A). Additionally, J-PKAcα in the chimeric holoenzyme is superimposable to the previously reported (Cheung et al., 2015) structure of the PKI-bound chimera with a Cα RMSD of 0.39 Å (Figure S5B). The fused J-domain is similarly tucked underneath the C-lobe, and the contact area for the J-domain in the chimera is ~380 Å^2^. The J-domain in the chimeric holoenzyme has significantly higher temperature factors (B factor) than the rest of the holoenzyme, even at this medium resolution, suggesting that it retains a high degree of flexibility in the holoenzyme, similar to its PKI-bound state in solution based on NMR experiments (Tomasini et al., 2018) (Figure 1C and Table S1). The heterodimer in the chimeric holoenzyme is also structurally similar to the previously solved R: PKAcα heterodimers (Figure S5C) (Taylor et al., 2012), showing the J-domain fusion to the PKAcα does not alter the PKAcα interactions with the RIα subunit.. The J-domains in the holoenzyme locate close to the CNB-B domain of the adjacent RIα subunit, with the shortest Cα atoms distance at ~8 Å (Figure 1D). Residues 1-91 of RIα are missing in the electron density although by SDS-PAGE and silver staining, we validated that full-length RIα and J-PKAcα are present in the protein crystal (Figure S3). This absence of electron density for the D/D domain and part of the following N-linker is likely related to the flexible nature of this region (Li et al., 2000).

**Table 1.** Data Collection and Refinement Statistics

| | RI$\alpha_2$:J-PKAc$\alpha_2$ | RI$\alpha_2$:PKAc$\alpha_2$ |
| --- | --- | --- |
| **Data collection** |  |  |
| Space group | P6<sub>5</sub>22 | P2<sub>1</sub>2<sub>1</sub>2<sub>1</sub> |
| No. of molecules in one asymmetric unit | 1 | 2 |
| Cell dimensions |  |  |
| *a*, *b*, *c* (Å) | 166.50, 166.50, 332.70 | 140.50, 186.16, 186.67 |
| $\alpha$, $\beta$, $\gamma$ (°) | 90, 90, 120 | 90, 90, 90 |
| Resolution (Å) | 50-3.66 (3.79-3.66) | 50-4.75 (4.92-4.75) |
| *R*<sub>sym</sub> (%) | 13.5 (49.8) | 10.9 (41.2) |
| *I* / σ*I* | 29.5 (8.7) | 16.3 (2.7) |
| Completeness (%) | 100.0 (100.0) | 97.2 (80.5) |
| Redundancy | 21.3 (22.2) | 6.7 (4.6) |
| **Refinement** |  |  |
| Resolution (Å) | 50-3.66 | 50-4.75 |
| No. reflections | 30810 | 24239 |
| *R*<sub>work</sub> / *R*<sub>free</sub><sup>a</sup> (%) | 20.0/25.0 | 21.2/25.5 |
| No. atoms |  |  |
| Protein | 11292 | 20336 |
| Ligand/ion | 66 | None |
| Water | None | None |
| *B*-factors |  |  |
| Protein | 110.69 | 269.92 |
| Ligand/ion | 91.06 | None |
| R.m.s deviations |  |  |
| Bond lengths (Å) | 0.003 | 0.014 |
| Bond angles (°) | 0.623 | 1.570 |

Values in parentheses are for the highest resolution shell.
<sup>a</sup> *R*<sub>free</sub> was calculated by using a 5% of randomly selected reflections.

### Highly dynamic J-domains in the chimeric fusion RIα_2_:J-PKAcα_2_ holoenzyme

Small Angle X-ray Scattering (SAXS) results (Figure S6 A-D) are in general consistent with the observed shape of the chimeric holoenzyme in solution. The calculated solution scattering data from the crystal structure fit to the SAXS solution experimental data reasonably well with a χ^2^ of 1.44. The chimeric holoenzyme in solution displays larger R_g_ and D_max_ values (Figure S6D, Table S2) than those calculated for the crystal structure. The dynamic D/D domain with the N-linker regions or the dynamic nature of the J-domain may account for the observed larger dimension of the chimeric RIα holoenzyme in SAXS experiments compared to the crystal structure (Figure S6D).

Previous MD simulations of isolated J-PKAcα (Tomasini et al., 2018) identified two representative conformational states (Figure 2A). In the highest occupied state, the J-domain was positioned beneath the C-lobe of the kinase core in a J-in state, which is similar to what we observe in our chimeric holoenzyme structure. A second state showed the J-domain rotated away from the core to form an extended J-out conformation, and this flexibility of the J-domain was confirmed by NMR studies (Tomasini et al., 2018). To probe the possible motions of the J-domain in the chimeric holoenzyme, we performed three 1 μs MD simulations of the chimeric holoenzyme starting from either the crystal structure, or from a conformation with the J-domain modeled onto the holoenzyme crystal structure in the J-in or J-out state. The simulations from the chimeric crystal structure showed the majority of conformations in an extended J-out state, and far from the RIα subunit (Figure 2B). This is in contrast to the simulation performed with free J-PKAcα (Tomasini et al., 2018), where the J-in state was the highest occupied state. In the simulation started from the J-in state model, the J-domain from one chimera rotated to an extended conformation while that of the other remained in a J-in state (Figure S7A) to form stable interactions with its adjacent R subunit (Figure S7B). The minimum distances between Cα atoms in the J-domain to any Cα atom in the adjacent RIα subunit over all simulations ranged from 5.1 Å to 31.2 Å, emphasizing the flexibility of the J-domain (Figure 2C). In the simulation starting from the J-out state model, the J-domains of both chimeric subunits remained in the J-out state throughout the 1 µs simulation and did not show any interaction with the R subunits (Figure S8). The calculated data from the three final MD simulation conformations of the chimeric holoenzyme (Figure S9 and Table S2), with one copy or both of the J-domain sampling the “out” state, are generally in agreement with the experimentally obtained SAXS solution data despite the lack of electron density for the D/D domain. The R_g_ and D_max_ values of these three MD simulation conformations of the chimeric holoenzyme are also closer to the SAXS solution data than that of the crystal structure, suggesting the extended J-out state is a likely conformation of the J-domain in the chimeric holoenzyme in solution.

**Figure 2.**
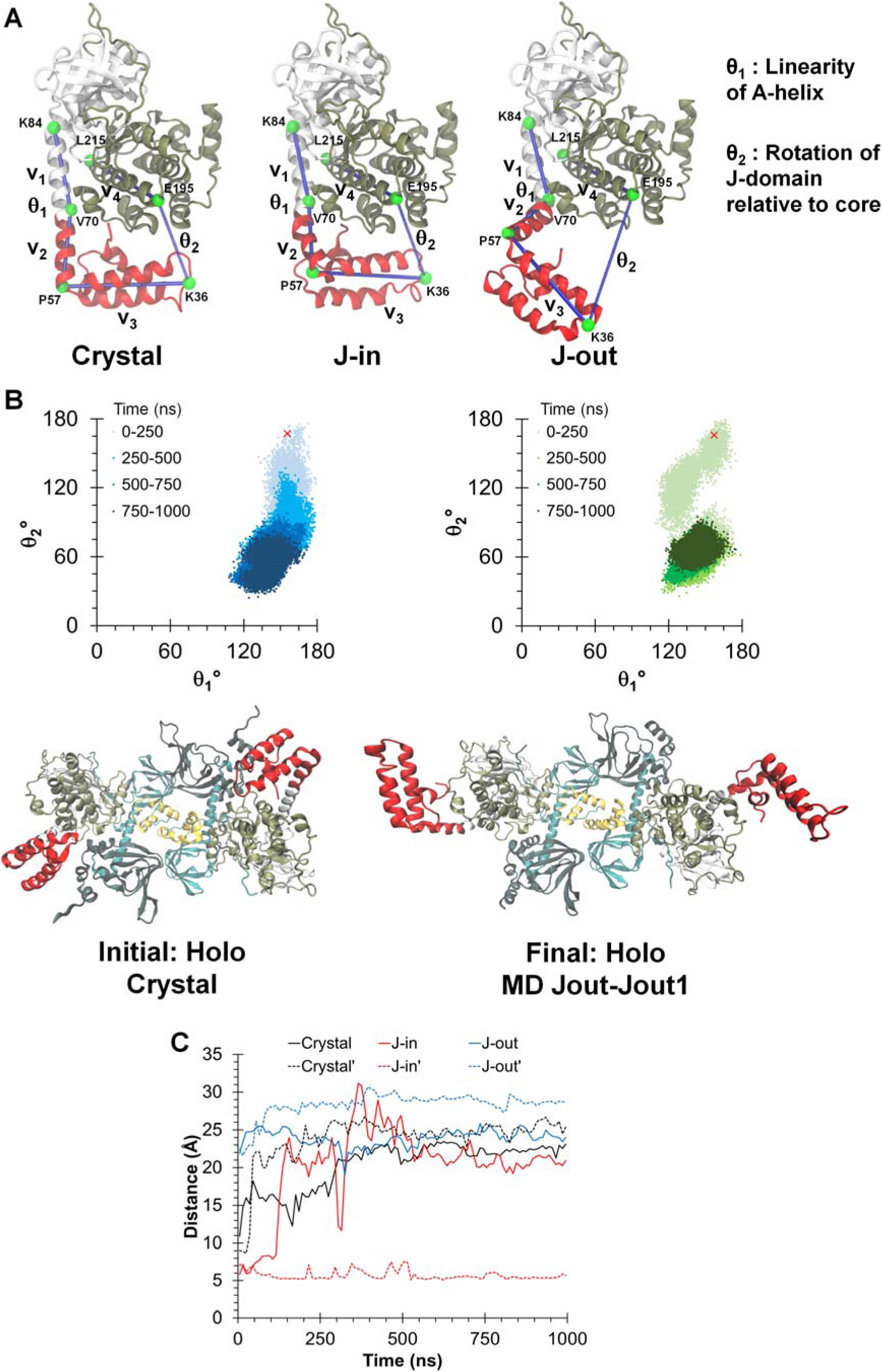
Dynamic conformations of J-domains in the chimeric holoenzyme during MD simulations. (A) Three different simulations were initiated from: the chimeric crystal structure (crystal), the J-in and J-out states. θ_1_ and θ_2_ are angles defined to probe the dynamics of the J-domain. (B) Top: Orientation of the J-domain for both copies of the chimera in the RIα_2_:J-PKAcα_2_ holoenzyme, as given by θ_1_ and θ_2_ over a 1 μs simulation of the chimeric holoenzyme starting from the crystal structure. The red ‘x’ indicates the position of the J-domain at the beginning of the simulation. Darker colors indicate later in time. Larger values of θ_2_ indicate a conformation in which the J-domain is tucked underneath the PKAcα core while smaller values indicate an extended conformation. Bottom: Initial and final conformations of the RIα_2_:J-PKAcα_2_ simulation started from the crystal structure. (C) Minimum Cα distances between the J-domain and the adjacent RIα subunit for the three simulations. Solid and dotted lines indicate each copy of the J-domain in the holoenzyme respectively.

### Isoform-specific interface between the RIα:J-PKAcα heterodimers

The interface between the chimeric heterodimers is solely created by the antiparallel alignment of the CNB-A and CNB-B domains in the RIα dimer, with a contact area of ~970 Å^2^(Figure 3A). Wedged against each other from the two-fold symmetry-related RIα subunits are the two CNB-A N3A motifs (Figure S10) which include the αN and αA helices as well as the connecting 3_10_-loop. A similar N3A-N3A’ interface was first reported in the cAMP-bound RIα homodimeric RIα_2_(cAMP)_4_ structure (Figure 3B) (Bruystens et al., 2014). Each αA-helix is perpendicular to the opposing αN’-helix, thus creating a rectangular shaped four-helical bundle interface. The RIα-RIα’ interface also contains two identical salt bridge contacts between E179 in the RIα CNB-A domain and R315’ and R340’ in the RIα’ CNB-B’ domain (Figure 3A). Similar to the cAMP-bound RIα dimer (Bruystens et al., 2014), the N3A-N3A’ helical bundle is mostly hydrophobic, involving residues M123, Y120 and F148 from each N3A motif. These hydrophobic interactions are generally stable throughout the course of the MD simulations. The helical bundle with its two-fold symmetry also includes two identical hydrogen bond networks. Residues Y120 and K121 in the αN-helix form hydrogen bonds with N142’, S145’ and D149’ in the αA’-helix. While not directly involved in interactions at the N3A-N3A’ interface, R144 in the αA-helix forms hydrogen bonds with the backbone oxygens of F136 and L139 from the 3_10_-loop. These hydrogen bonds break during cAMP activation as a consequence of outward motion of the 3_10_-loop. Mutations of residues R144 and S145 are associated with CNC disease, which creates a holoenzyme that is poorly regulated and more easily activated by cAMP (Park et al., 2012). Substitutions of R144, S145 and N3A interface residues Y120 and F148 caused increased sensitivity for cAMP activation of the corresponding RIα holoenzymes and reduced cooperativity for cAMP binding (Bruystens et al., 2014). The Hill Coefficient for R144S and S145G were reduced to 1.4-1.5 while the Hill Coefficient was 1.0-1.1 for the Y120A and F148A mutants. The N3A-N3A’ helical bundle was also seen in truncated RIα monomer structures as an interaction site for crystal packing (Badireddy et al., 2011; Wu et al., 2004a, 2004b). However, this interface is not observed in any structures associated with RIIα or RIIβ. Sequence alignment also shows that RII subunits lack most of the key residues involved in forming the N3A-N3A’ interface (Figure 3A), emphasizing again that the N3A-N3A’ four-helical bundle is isoform-specific.

**Figure 3.**
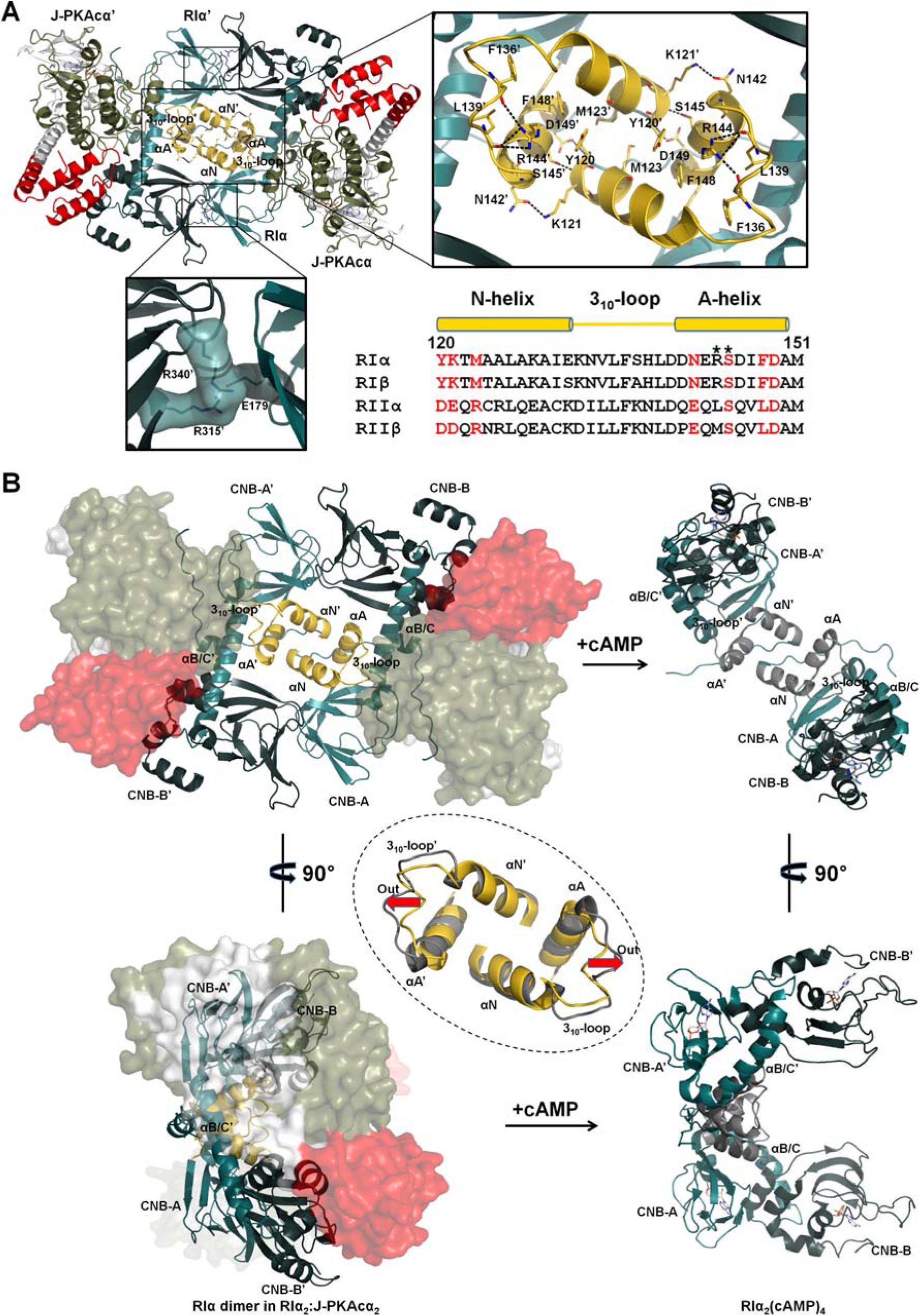
Interactions of the two RIα:J-PKAcα heterodimers in the chimeric holoenzyme. (A) Overall interface of the two heterodimers consists of a large N3A-N3A’ interface and two identical small interfaces with salt bridges. Sequence alignment of the N3A motifs from different R isoforms is shown at the right bottom. Interface residues at the N3A motif are labeled in red. CNC mutations are marked with asterisks. (B) The N3A-N3A’ four-helical bundle acts as a structural anchor during cAMP activation. The αB/C-helix and the CNB-B domain of RIα undergo dramatic conformational changes upon cAMP binding, while the N3A-N3A’ helical bundle is almost unaltered except the move-out of the 3_10_-loops shown by the red arrows. The superimposition of the N3A-N3A’ interfaces in the chimeric holoenzyme and the cAMP-bound RIα dimer (gray, PDB ID 4MX3) is shown in the dashed circle.

The overall structure of the N3A-N3A’ helical bundle is also conserved in the cAMP-bound RIα homodimer (Figure 3B). Thus, the N3A-N3A’ bundle likely serves as a structural anchor and contributes to the activation of the holoenzyme by cAMP activation and the following dissociation of R and C subunits. By contrast, the extended αB/C-helix that connects the CNB-A and CNB-B domains in the holoenzyme adopts a bent configuration in the cAMP-bound RIα homodimer with the CNB-B domain rotated dramatically to a position underneath the relatively stable CNB-A domain (Figure 3B). Moreover, the R315’-E179-R340’ salt bridge interactions observed in the holoenzyme (Figure 3A) between the RIα dimer become broken in the cAMP-bound RIα homodimer.

### Overall structure of wt RIα_2_:PKAcα_2_ demonstrates two distinct holoenzyme conformations

To determine if the structure of the chimeric holoenzyme is unique to the fusion chimera protein, the wt RIα holoenzyme was formed *in vitro* by mixing the individually purified subunits followed by gel filtration, and its structure was determined at 4.75 Å resolution. It required different crystallization conditions (Figure 4 and Table 1) and has a distinct space group (P2_1_2_1_2_1_) compared to the chimeric RIα_2_:J-PKAcα_2_ holoenzyme and the previously the solved structures of the wt tetrameric RIβ_2_:PKAcα_2_ or RIIβ:PKAcα_2_ holoenzymes (Ilouz et al., Zhang et al., 2012). Each ASU contains four RIα:PKAcα heterodimers. The presence of full-length proteins in the crystals was confirmed by SDS gel analysis and silver staining, as described earlier (Figure S11). Analysis of the crystal packing showed that each ASU has two RIα_2_:PKAcα_2_ holoenzyme molecules with distinct quaternary structures. While holoenzyme 1 has a conformation almost identical to the chimeric holoenzyme (Figures. 4A, S12A and S13A), holoenzyme 2 has a much smaller N3A-N3A’ interface with an area of ~370 Å^2^ created only by the αN-helices (Figure 4B and S13B) and thus has a conformation distinct from holoenzyme 1. Both of the holoenzyme molecules contain two RIα:PKAcα heterodimers with a rotational two-fold symmetry through the N3A-N3A’ interface. The heterodimers of RIα:PKAcα are almost identical in the two different tetrameric holoenzyme conformations (Figure S12B) with a Cα RMSD of 0.26 Å and also resemble the previously published structure of a truncated RIα(91-379):PKAcα heterodimer (Figure S12C) with a Cα RMSD of 0.86 Å (Kim et al., 2007). Similar to that in the chimeric holoenzyme, in both of the wt conformations, CNB-A is juxta-positioned against CNB-B’ thus supporting the enhanced allostery that is associated with the RIα_2_:PKAcα_2_ holoenzyme compared to the RIα:PKAcα heterodimer (Taylor et al., 2012). RIα competition assay results showed that the chimera and wt PKAcα have similar ability for RIα association (Figure 4C). In addition, the chimeric and wt RIα holoenzymes have no significant differences in cAMP activation nor its cooperativity (Figure 4C).

**Figure 4.**
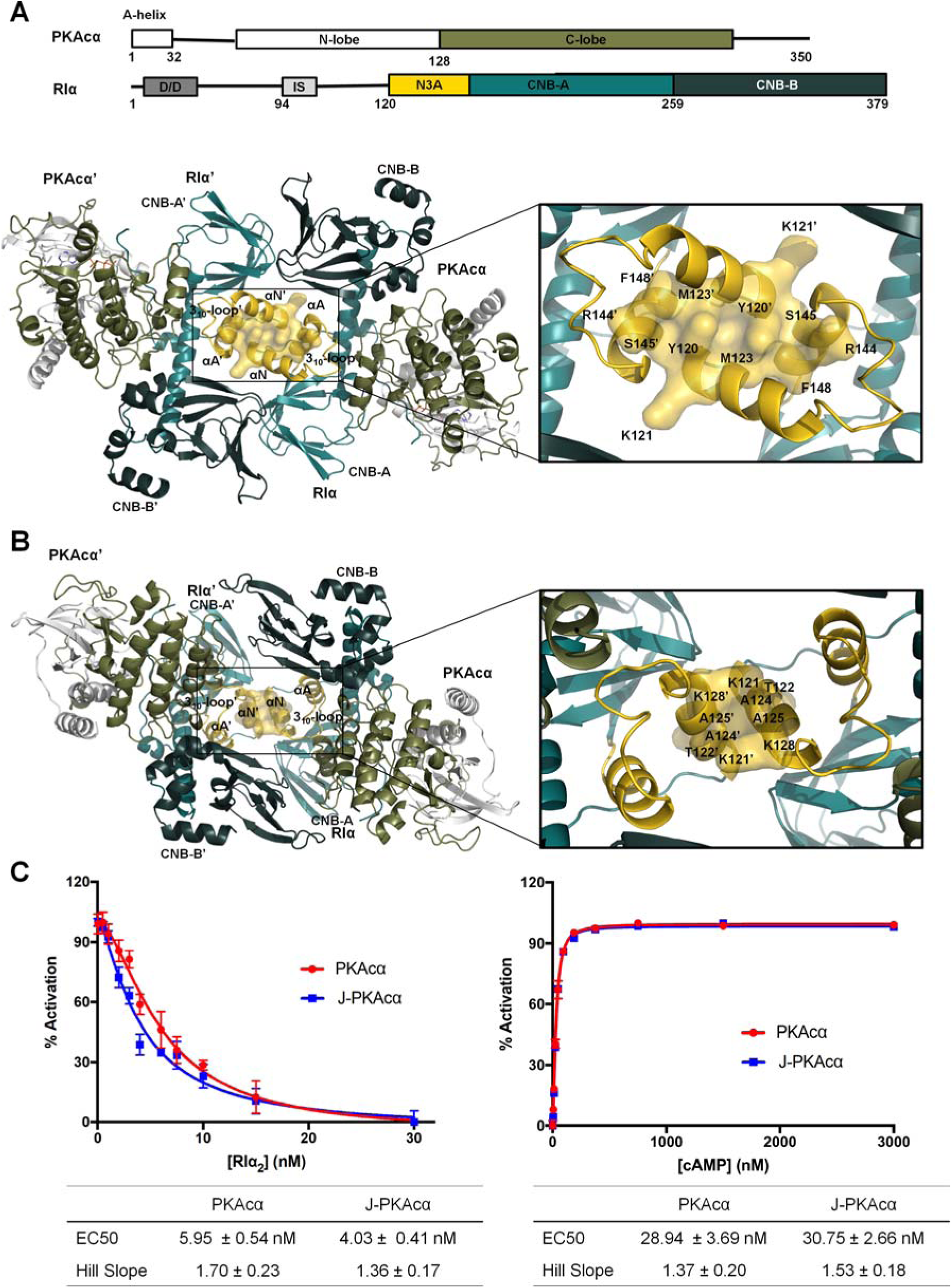
Interactions of the two RIα:PKAcα heterodimers in the wt holoenzyme. (A) Interface of the two heterodimers in the wt holoenzyme 1. Domain organization and color coding of the PKAcα and RIα subunits are shown on the top. (B) Interface of the two heterodimers in the wt holoenzyme 2. (C) Fluorescence polarization assays to measure RIα inhibition (left) by PKAcα (red) and J-PKAcα (blue) as well as holoenzyme activation by cAMP (right). All data points are mean ± s.d. (n = 3 independent experiments).

The PKAcα subunits in holoenzyme 2 become closer to the heterodimer interface and to the symmetry-related RIα subunit than in holoenzyme 1 (Figure S12D). During the MD simulation, PKAcα in holoenzyme 2 is capable of interacting further with RIα’ (Figure S12E). MD simulations of each of the two conformations of the wt RIα_2_:PKAcα_2_ holoenzyme indicates that over the 1 μs of the simulation they are stable and do not interconvert (Figure S14A). The interfacial area between the RIα dimer in the wt holoenzyme 1 crystal structure resembles that of the chimeric holoenzyme simulations. The contact area in holoenzyme 2 is slightly increased (Figure S14B), which is largely due to a slight rotation of the RIα-RIα’ interface during the MD simulation.

### Isoform-specific quaternary structures of PKA holoenzymes

The chimeric and wt RIα holoenzymes have quaternary structures different from the previously solved wt RIβ and RIIβ holoenzymes, even though the structures of all PKA heterodimers are remarkably similar (Figure 5). The quaternary structure isoform diversity is essential for each holoenzyme to create a distinct signaling hub that can respond to local levels of second messengers such as cAMP, and allows formation of distinct macromolecular complexes with local substrates and accessory proteins at different cellular sites.

**Figure 5.**
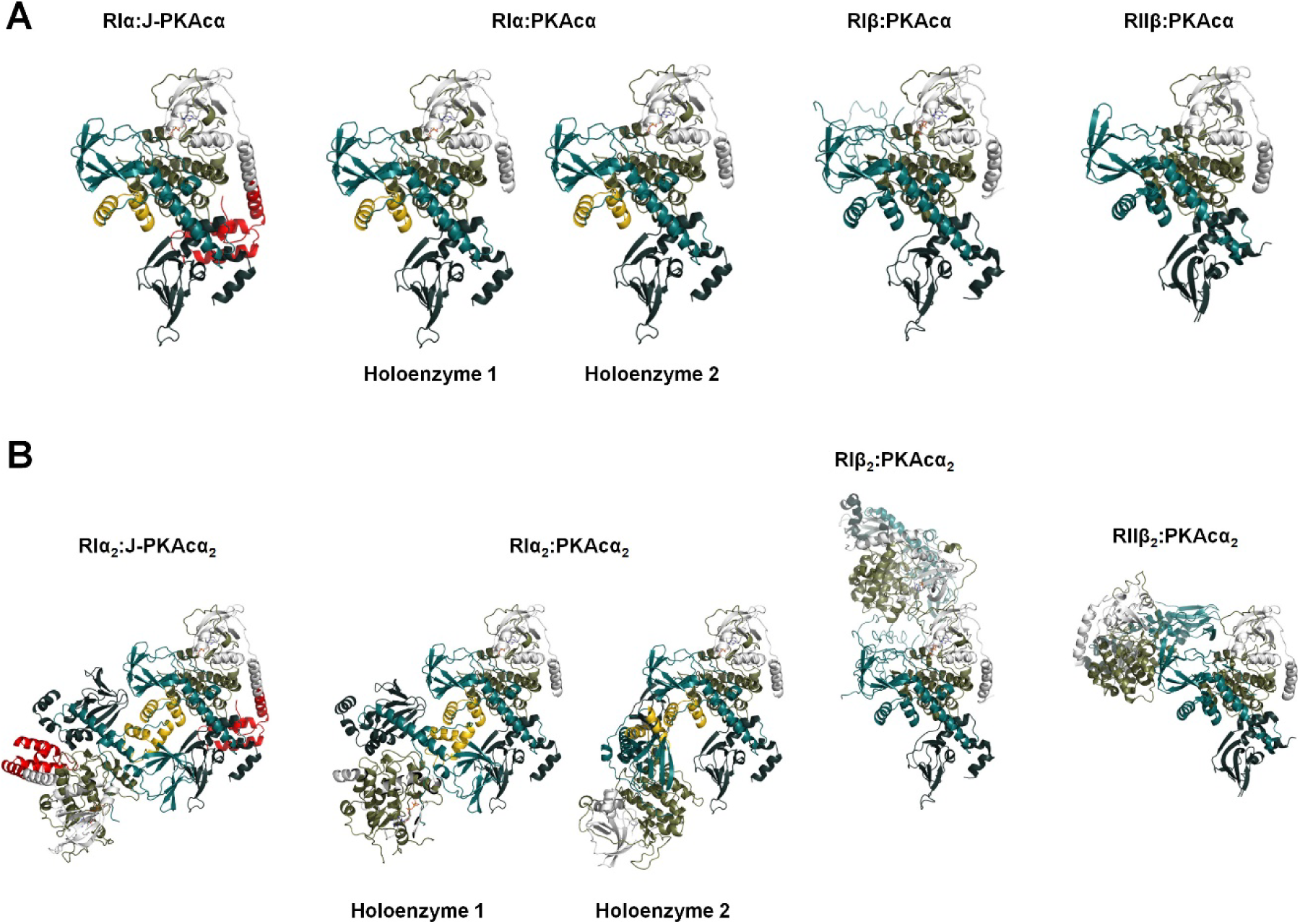
Structural comparison of PKA holoenzymes. (A) Side-by-side view of heterodimers at the same orientation: structures of RIα:J-PKAcα, RIα:PKAcα, RIβ:PKAcα (PDB ID 4DIN) and RIIβ:PKAcα (PDB ID 3TNP) heterodimers in the respective PKA holoenzymes. (B) Structure comparison of the chimeric RIα_2_:J-PKAcα_2_ and wt RIα, RIβ and RIIβ holoenzymes.

The RIα holoenzymes and the view of CNB-B movement in these holoenzymes reported in this study (Figures. 1 and 4) are distinct from the earlier model of the RIα holoenzyme that was based on crystal packing of two truncated RIα(73-244):PKAcα heterodimers (Boettcher et al., 2011). The earlier RIα holoenzyme model showed the two R:PKAcα heterodimers have cross talks between the CNB-A domain of one R: PKAcα dimer with the PKAcα’ of the other dimer and also allowed the modeled-in CNB-B domain movement. Such mobility of the CNB-B domain is consistent with previously obtained SAXS data of the RIα(91-379):PKAcα heterodimer and led to a suggestion that the CNB-B domain of RIα is mobile and moves away from PKAcα with Gly235 serving as a hinge point (Cheng et al., 2009). Recent studies have shown that CNB-B domain flexibility is linked to cAMP activation in the RIα(91-379):PKAcα truncated heterodimer (Hirakis et al., 2017; Barros et al., 2017). However, this view of CNB-B movement in the holoenzyme is different from the packing observed here in the full-length chimeric and wt RIα holoenzymes (Figures. 1 and 4) where the CNB-B domain interacts with the opposite CNB-A’ domain; this interaction would prevent the suggested hinge motion in the holoenzyme. Consistent with our full-length RIα holoenzyme structure, MD simulations monitoring the dynamics of the αB/C-helix indicate it to be stable in the full-length holoenzyme with a near linear average of ~162° in all simulations (Figure S15).

### Effects of the J-domain on PKAcα function

The discovery that J-PKAcα is an oncogenic driver of FLHCC and thus a therapeutic target represents a significant breakthrough for FLHCC research (Honeyman et al., 2014). The fusion of the J-domain to PKAcα (Honeyman et al., 2014; Kastenhuber et al., 2017) may lead to alterations in kinase activity, substrates, dynamics, location or regulation at the level of the kinase subunit, holoenzyme and/or even higher molecular complexity level. As shown in the RIα competition as well as the cAMP activation assays, no significant differences were observed in terms of RIα association with either the chimera or wt PKAcα (Figure 4C) and the addition of the J-domain does not impact the sensitivity of the chimeric holoenzyme to cAMP activation (Figure 4C). As suggested by a thermostability assay (Figure 6A), the dynamic J-domain does not introduce a significant destabilizing effect on the chimera, nor on the chimeric RIα holoenzyme (Table S3). Similarly, J-PKAcα displayed unaltered binding affinities for ATP and inhibitor peptide (Figure 6B). Additionally, in agreement with previous reports (Cheung et al., 2015), the chimeric protein was slightly more active than its wt counterpart with unchanged enzymatic efficiency as shown by *k*_cat_/K_m_ values (Figure 6C), suggesting that the J-domain may affect PKAcα enzyme dynamics. In the crystal structure of the chimeric fusion RIα_2_:J-PKAcα_2_ holoenzyme, the presence of the J-domain does not prevent formation of the holoenzymes, and the C-subunit, where the J-domain fusion occurs, is not at the symmetrical interface in the holoenzyme between the two RIα:J-PKAcα heterodimers (Figure 1). Thus, rather than affecting the PKAcα interactions with the regulatory subunits, it is possible that addition of the J-domain alters the conformational landscape of the chimeric fusion holoenzymes, impacting interactions with other molecules. The higher B-factors in the J-domain suggested a large degree of conformational flexibility (Figure 1C, Table S1). MD simulations indicate a wide range in the conformational diversity of the J-domain appendage both in isolated J-PKAcα and in the holoenzyme, perhaps influencing enzyme dynamics through allosteric networks or holoenzyme interaction with other proteins.

**Figure 6.**
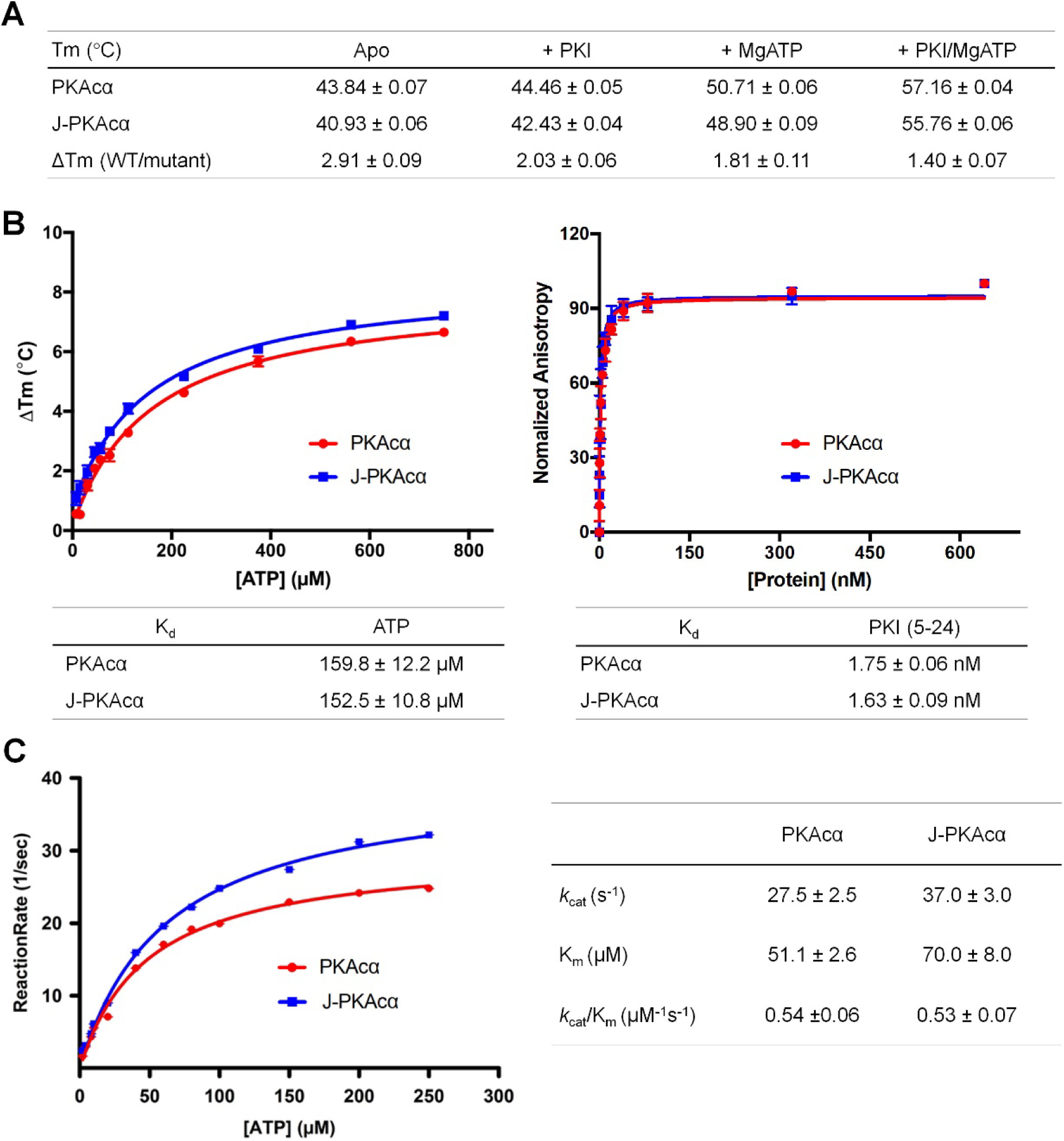
Stability, ATP binding and kinetic studies of J-PKAcα and PKAcα. (A) Stability of J-PKAcα and PKAcα (apo, with ATP, and/or with PKI binding) measured by thermofluor assay. (B) ATP and PKI binding affinities of J-PKAcα (blue) and wt PKAcα (red). ATP binding (left) and PKI binding (right) were measured by thermofluor and fluorescence anisotropy assay respectively. (C) Steady-state kinetics of phosphotransfer reaction of J-PKAcα (blue) and wt PKAcα (red). All data points are mean ± s.d. (n = 3 (panels a and b) and 2 (panel c) independent experiments).

## Discussion

The oncogenic J-PKAcα has been crystallized here for the first time in one of its most important physiological states where it is associated in a holoenzyme complex with the RIα subunit. This structure demonstrates that the N-terminal fusion does not interfere with the general organization of the R_2_:PKAcα_2_ holoenzyme, and this also has relevance for the various PKAcα isoforms some of which have large extensions at the N-terminus (Søberg et al., 2017). Comparing the conformational states of the wt and chimeric RIα holoenzymes that display some novel interfaces may guide the development of drugs that selectively target not only to the J-domain and catalytic core to directly block chimera activity, but also regions present only at the holoenzyme level to block holoenzyme activation. The presence of alternate conformations of the holoenzymes may constitute a way to target the chimera selectively, as the conserved activity site of the wt PKAcα and the chimeric fusion J-PKAcα have little structural differences and the enzyme function is barely affected by the J-domain fusion. The enhanced dynamics of the chimeric holoenzyme may also expose some sites that are otherwise too transient to target. It may also be possible to trap a dynamic state independent of whether the holoenzyme is dissociated or not. Using a strategy that simultaneously blocks the activity of the oncogenic driver kinase and/or its holoenzyme dissociation would significantly reduce the possibility that a random mutation in the driver enables the tumor cells to escape treatment. RIα is a critical master switch for regulating PKA activity in cells, and it is likely that unregulated PKA activity is important, at least in part, for driving FLHCC. The importance of RIα is further supported by the recent finding that in a few rare cases, CNC mutations in RIα can drive FLHCC. Most CNC mutations, including the haplo-insufficiency caused by nonsense mediated decay of the RIα messenger RNA, do not drive FLHCC, so the unregulated phenotype associated with CNC is not in itself sufficient to explain these rare CNC mutations that are associated with FLHCC.

MD simulations show that the J-domain is highly dynamic in the chimeric RIα holoenzyme. The presence of the J-domain will likely also alter the phosphor-proteome of the tumor cells. At this point it is not clear how the presence of the J-domain influences the function of the PKA holoenzymes in cells. The wt PKAcα is also myristylated at its N-terminus and we have shown previously that this can be important for targeting the RIIβ holoenzymes to membranes (Zhang et al., 2015). This acylation site is missing in the fusion chimera protein and may also contribute to dysfunctional PKA signaling. Interestingly, the striking similarity on the overall structures and biochemical properties of the wt and chimeric RIα holoenzymes suggests the specificity of chimeric holoenzyme in its role in FLHCC need to be further sought at another level. It will be extremely important to elucidate how the conformational state and abundance of the different holoenzymes in the tumor cells and the holoenzymes communicate with their neighbors and substrates. In particular, it is important to determine how these macromolecular assemblies are altered in FLHCC by comparing paired tumor and adjacent normal liver samples. Understanding in detail how J-PKAcα signaling pathways drive disease will shed light on understanding its transformation to FLHCC and is expected to improve diagnosis and therapeutic treatment for this cancer.

## Methods

### Protein expression, purification and crystallization

Bovine wt full-length RIα was expressed in *Escherichia coli* (*E. coli*) BL21 (DE3) pLysS and purified as described before (Barros et al., 2017) Both human full-length J-PKAcα and PKAcα were engineered with an N-terminal His_6_SUMO tag. The constructs were then transformed into *E. coli* BL21 (DE3) for protein expression. The starter cultures were grown in LB media with 50 µg/mL kanamycin overnight at 37 °C and then 1:100 diluted into the same media. The cultures were grown at 37 °C until the cell density reached 0.5-0.6 OD_600_, after which the temperature was lowered to 24 °C and protein expression was induced overnight by adding isopropyl β-D-thiogalactoside to a final concentration of 0.5 mM. Cells were harvested by centrifugation, resuspended in lysis buffer containing 20 mM Tris-HCl pH 8.0, 100 mM NaCl, 5 mM β-mercaptoethanol and lysed by microfluidizer. The lysates were centrifuged, and collected supernatants were incubated with Ni-nitrilotriacetic (Ni-NTA) agarose beads overnight at 4 °C. The beads were rinsed with lysis buffer and then 10X bed volume of wash buffer (lysis buffer plus 20 mM imidazole). The proteins were eluted with 3X bed volume of elution buffer 1 (lysis buffer plus 50 mM imidazole) and elution buffer 2 (lysis buffer plus 100 mM imidazole). The eluates were spin dialyzed into the lysis buffer, after which NP-40 was added to them to a final concentration of 0.1%, and subjected to U1P1 (an engineered SUMO protease) digestion for 1 h at 25 °C at a molar ratio of 200:1 (protein:enzyme) to remove the His_6_SUMO tag. The cleaved tag and the protease were then removed from the proteins using Ni-NTA beads. Then the full-length RIα_2_:J-PKAcα_2_ and RIα_2_:PKAcα_2_ holoenzymes were formed by mixing RIα with J-PKAcα or PKAcα in a 1:1.5 molar ratio and spin dialyzed into a holoenzyme buffer containing 50 mM MOPS pH 7.0, 50 mM NaCl, 1 mM TCEP, 1 mM MgCl_2_ and 0.1 mM ATP. The formed complexes were loaded onto Hiload 16/600 Superdex 200 pg size exclusion column preequilibrated with the same buffer. Proteins from the peak fractions corresponding to the holoenzymes were collected, concentrated to ~10 mg/mL and subjected to extensive crystallization screening or used for biochemical assays. Crystallization was conducted at 20 °C using the hanging drop vapor diffusion method by mixing the protein and precipitants at a ratio of 1:1. The RIα_2_:J-PKAcα_2_ crystals were grown in a buffer containing 100 mM NaCl, 16-18% pentaerythritol propoxylate and 10% dimethyl sulfoxide and to their final size in ~2 weeks. The RIα_2_:PKAcα_2_ crystals were grown in a buffer containing 100 mM HEPES sodium-MOPS (acid) pH 7.5, 90 mM NPS (30 mM sodium nitrate, 30 mM sodium phosphate dibasic, 30 mM ammonium sulfate), 40-42% Precipitant Mix 2 (40% ethylene glycol; 20% PEG 8000), 3% D-(+)-glucose monohydrate and to their final size in ~3 weeks.

### Structure determination

Diffraction data were collected at the 22ID beamline of the Advanced Photon Source (APS), Argonne National Laboratory (ANL). Data were indexed, integrated and scaled using the HKL2000 program (Otwinowski et al., 1997). The best RIα_2_:J-PKAcα_2_ and RIα_2_:PKAcα_2_ holoenzyme crystals diffracted to 3.66 and 4.75 Å, respectively. The initial phase of RIα_2_:J-PKAcα_2_ was determined using program PHASER (McCoy et al., 2007) with the structures of PKAcα Δexon1 (from PDB ID 4WB8) (Cheung et al., 2015) and RIα (from PDB ID 2QCS) (Barros et al., 2017) as search models. Refinement of the molecular replacement model was performed with PHENIX (Adams et al., 2010) and COOT (Emsley et al., 2004) alternatively. Initially, three rounds of Cartesian, individual B-factors, atomic occupancies and Cartesian simulated annealing (start temperature 5,000 K) refinement were performed in PHENIX, with the restraints of torsion-angle non-crystallographic symmetry (NCS), reference models and secondary structures. The reference models were J-PKAcα (from PDB ID 4WB7) (Cheung et al., 2015) and RIα (from PDB ID 2QCS) (Barros et al., 2017). In addition, stereochemistry and atomic displacement parameters weights were optimized during the refinement. The final refinement protocol included three rounds of Cartesian, individual B-factors and atomic occupancies refinement. The final RIα_2_:J-PKAcα_2_ model has 92.5% of residues in the favored Ramachandran region and 7.5% in the allowed region. The initial phase of RIα_2_:PKAcα_2_ was determined using program PHASER with the refined structure of the J-domain omitted RIα:J-PKAcα as the search model. Refinement of the molecular replacement model was carried out with REFMAC5 (Nicholls et al., 2012), PHENIX and COOT. First, rigid body refinement was performed using REFMAC5. Then 10 rounds of Cartesian, group B-factors (single residues were divided into mainchain and sidechain), atomic occupancies and Cartesian simulated annealing (start temperature 5,000 K) refinement were performed in PHENIX, with the restraints of global NCS, reference models (from PDB ID 2QCS) and secondary structures. The final refinement protocol included three rounds of Cartesian, individual B-factors and atomic occupancies refinement with the global NCS restraint. The final RIα_2_:PKAcα_2_ model has 84.6% of residues in the favored Ramachandran region and 15.4% in the allowed region. Data collection and refinement statics are summarized in Table 1. Models were evaluated using the MolProbity web server (molprobity.biochem.duke.edu/).

### Small angle X-ray scattering (SAXS) experiment

SAXS measurements were performed at the 12ID-B beamline of APS, ANL. Photon energy was 13.3 KeV, and sample-to-detector distance was 3.6 m. To minimize radiation damage, thirty image frames were recorded with an exposure time of 1-2 s for each buffer and sample solution using a flow cell. The 2D images were reduced to 1D scattering profiles, and then grouped by sample and averaged using the MatLab software package at the beamlines. Concentration series measurements for the same sample were carried out to remove the scattering contribution due to interparticle interactions and to extrapolate the data to infinite dilution. The concentrations were 0.5, 0.7 and 0.9 mg/ml for RIα_2_:J-PKAcα_2_ in the buffer containing 50 mM MOPS pH 7.0, 50 mM NaCl, 1 mM TCEP, 1 mM MgCl_2_ and 0.1 mM ATP. The buffer background subtraction and intensity extrapolation to infinite dilution were carried out using NCI in-house developed MatLab script NCI-SAXS. Theoretical scattering profiles were generated from crystal structure and models and compared with the experimental SAXS data at q < 0.5 Å^-1^ using the CRYSOL software (Svergun et al., 1995). The pair-distance distribution function P(r) and maximum dimension (D_max_) were generated using GNOM (Svergun et al., 1992).

### Kinase activity assay

The enzymatic activity of wt PKAcα or J-PKAcα was measured spectrophotometrically with a coupled enzyme assay (Cook et al., 1982). The ADP formation is coupled to the pyruvate kinase (PK) and lactate dehydrogenase (LDH) reactions. The reaction rate is determined by following the decrease in absorbance at 340 nm at 25 ͦC on a Photodiode Array Lambda 465 UV/Vis Spectrophotometer (PerkinElemer). The Michaelis-Menten parameters for ATP were determined by fixing Kemptide substrate (LRRASLG) at saturating concentrations while varying the concentrations of ATP. Reactions were pre-equilibrated at room temperature and initiated by adding ATP. The kinase reaction mixture contained 100 mM MOPS pH 7.1, 50 mM KCl, 6 mM phosphoenolpyruvate, 0.5 mM nicotinamide adenine dinucleotide (NADH), 100 µM of Kemptide, 15 units of LDH, 7 units of PK, and varying concentrations of ATP from 0 to 250 µM. MgCl_2_ was present in a constant 1 mM excess over ATP. The data was analyzed and fitted to the Michaelis-Menten equation using SigmaPlot software.

### Inhibitor peptide PKI binding assay

Fluorescence anisotropy was used to measure PKI to PKAcα or J-PKAcα. 0.9 nM FAM-labeled PKI (5-24) peptide was mixed with 0-2000 nM PKAcα or J-PKAcα in buffer containing 20 mM MOPS pH 7.0, 150 mM NaCl, 10 mM MgCl_2_, 1 mM ATP, and 0.01% Triton X-100. Fluorescence anisotropy was measured by using GENios Pro micro-plate reader (Tecan) in black flat-bottom costar assay plates with 485 nm excitation and 535 nm emission. The data was analyzed and fitted to the anisotropy single association hyperbolic equation using Prism software.

### RIα competition assay for catalytic subunit binding

Fluorescence polarization assay was used to measure the competition of RIα subunit with IP20 for wt PKAcα or J-PKAcα. 2 nM N-terminus FAM-labeled PKI peptide (5-24), and 10 nM PKAcα or J-PKAcα were mixed in the buffer containing 20 mM HEPES pH 7.0, 75 mM KCl, 0.005% Triton X-100, 10 mM MgCl_2_, 1 mM ATP, and 1 mM DTT. Two-fold serial dilutions of RIα from 30 nM to 0 nM were added to the PKI-bound catalytic subunits, followed by fluorescence polarization measurements using GENios Pro micro-plate reader (Tecan) in black flat-bottom costar assay plates with 485 nm excitation and 535 nm emission. The data was analyzed and fitted to the EC50 dose-response equation using Prism software.

### Stability assay

ThermoFluor assay was used to measure the stabilities of apo PKAcα or J-PKAcα subunits and its ATP and/or peptide binding forms. The reaction was conducted with 5 μM of proteins in 45 μL of the buffer containing 20 mM MOPS pH 7.0, 150 mM NaCl. Ligands were used at the following concentrations 1 mM ATP, 10 mM MgCl2, and 25 μM PKI peptide (5-24). For each ligand, triplicate reactions were measured in a 96-well plate. After proteins and ligands were mixed and incubated for 5 min on ice, 5 μL of 200X SYPRO Orange dye was added to each reaction. The samples were heated from 20 to 85 °C with a 0.5 °C/min heating rate by using CFX96 Real-Time PCR Detection System (Bio-Rad) in temperature scanning mode. The fluorescence signals were measured using the ROX channel.

### ATP binding assay

ATP dissociation constants were determined using the ThermoFluor assay. Similar condition as thermostability assay was used for ATP binding. The reactions were carried out in the buffer containing 20 mM MOPS pH 7.0, 150 mM NaCl with a range of ATP concentrations from 0 to 0.75 mM. After mixed with PKAcα or J-PKAcα, and incubated for 5 min on ice, 5 μL of 200X SYPRO Orange dye was added to each reaction. The samples were heated from 20 to 85 °C with a 0.5 °C/min heating rate by using CFX96 Real-Time PCR Detection System (Bio-Rad) in temperature scanning mode. The final concentration 4.5 μM of catalytic subunits was used to fit the data. The fluorescence signals were measured using the ROX channel. Each melting temperature was recorded and plotted versus ATP concentration.

### PKA cAMP activation assay

Fluorescence polarization assay was used to measure the activation of wt and chimeric RIα holoenzymes. 2 nM N-terminus FAM-labeled PKI peptide (5-24), 7.2 nM RIα_2_, and 12 nM catalytic subunit (wt or chimera) were mixed in the buffer containing 20 mM HEPES pH 7.0, 75 mM KCl, 0.005% Triton X-100, 10 mM MgCl_2_, 1 mM ATP and 1 mM DTT. To activate PKA catalytic subunits, 2-fold serial dilutions of cAMP from 3000 nM to 0 nM were added. The fluorescence polarization was measure by using GENios Pro micro-plate reader (Tecan) in black flat-bottom costar assay plates with 485 nm excitation and 535 nm emission. The data was analyzed and fitted to the EC50 dose-response equation using Prism software.

### Molecular dynamics simulations

MD simulations were performed to probe the dynamics of the RIα holoenzyme complexes. As previous simulations of the isolated J-PKAcα indicated a wide ensemble of conformations for the J-domain appendage (Tomasini et al., 2018), we performed three different simulations of RIα_2_:J-PKAcα_2_ with differing initial positions of the J-domain: the crystal structure, a J-in state model in which the J-domain was positioned close to the core of the catalytic subunit, and a J-out state model in which the J-domain was rotated away from the core of the catalytic subunit and the R:J-PKAcα interface. The J-domain conformations of the J-in and J-out states were those found in Tomasini *et al.* (Tomasini et al., 2018) as the top two representative conformations in a series of simulations performed on the isolated J-PKAcα. These two conformations of the J-domain were modeled onto the RIα_2_:J-PKAcα_2_ crystal structure. A similar methodology was used to model the first 14 amino acids and myristoylation motif which were missing from both conformations of RIα_2_:PKAcα_2_.

Structures were processed using the Protein Preparation Wizard in Maestro, solvated in a rectangular box with ~60,000 SPC waters and 150 mM sodium and chloride ions. Simulations were performed using the Desmond MD Package (Bowers et al., 2006) using the OPLS3 force field (Harder et al., 2016). Each system was subject to energy minimization using the steepest decent method succeeded by 100 ps of Brownian Dynamics simulation at constant volume and a temperature of 10 K with heavy atoms constrained. Subsequent equilibration included a 12 ps simulation at constant volume and at 10 K with heavy atoms restrained, followed by a 12 ps simulation at constant pressure with heavy atoms restrained, and finally a heating simulation in which the restraints were gradually relaxed and the system heated to 300 K over 24 ps. For production runs, the temperature was kept at 300 K using a Nose-Hoover Chain thermostat with a relaxation time of 1 ps (Martyna et al., 1992). The pressure was controlled at 1 bar using the Martyna-Tobias-Klein barostat with a relaxation time of 2 ps (Tuckerman et al., 2006). An integration time-step of 2 fs was used. Production simulations were performed for 1 μs saving system snapshots every 25 ps.

## Acknowledgments

We thank Drs. Alexandr Kornev, Di Xia, and Kylie Walters for critical reading of the manuscript and helpful discussions. We acknowledge use of the SAXS Core facility of Center for Cancer Research (CCR), National Cancer Institute (NCI) which is funded by FNLCR contract HHSN261200800001E and the intramural research program of the NIH, NCI, CCR. X-ray diffraction and SAXS data were collected at the 22ID and 12ID-B beamlines of the Advanced Photon Source, Argonne National Laboratory, respectively. We thank the Biophysics Resource in the Structural Biophysics Laboratory, NCI at Frederick for assistance. This work was supported by the National Institutes of Health grant GM34921 (S.S.T.), the Department of Defense grant CA160446 (S.M.S), and NIH grant 5R56CA207929 (S.M.S.), and the Intramural Research Program of the NIH, NCI, CCR (Zhang lab).

## Author contributions

B.C. carried out the protein purification, crystallization and structure determination work with help from J.M.F; TW.L. prepared plasmids and developed the protein purification protocol under the guidance of S.S.T. and P.Z; TW.L. and J.M.F did kinetic experiments; M.T. and S.M.S. performed the molecular dynamics simulation; L.F. and J.M.F. performed the SAXS experiments; B.C., J.M.F., TW.L., M.T. and P.Z. analyzed the data and wrote the paper with comments from all authors; P.Z. supervised all aspects of the project.

## Data availability

Coordinates and structure factors have been deposited in the Protein Data Bank with accession numbers 6BYR (RIα_2_:J-PKAcα_2_) and 6BYS (RIα_2_:PKAcα_2_).

## Additional information

Authors declare no competing interests.

---

**Table S1.** Averaged B factors for the J-domain and the rest of J-PKAcα bound with RIα or PKI

|  | B<sub>J</sub><sup>a</sup> | B<sub>C</sub><sup>b</sup> | Ratio of B<sub>J</sub>:B<sub>C</sub> |
| --- | --- | --- | --- |
| RIα<sub>2</sub>:J-PKAcα<sub>2</sub> | 203.45 | 94.09 | 2.16:1 |
| J-PKAcα:PKI<sup>c</sup> | 63.54 | 34.39 | 1.85:1 |

<sup>a</sup> B<sub>J</sub>: Averaged B factors of the J-domain (residues 1-69)
<sup>b</sup> B<sub>C</sub>: Averaged B factors of the rest of J-PKAcα (residues 70-405)
<sup>c</sup> from PDB ID 4WB7 (Cheung et al., 2015)

**Table S2.** SAXS structural parameters

| Holoenzyme conformation | $R_g$ (Å) in<br>reciprocal space | $R_g$ (Å) in<br>real space | $D_{\max}$ (Å) | $\chi^2$<sup>c</sup> |
| --- | --- | --- | --- | --- |
| SAXS solution scattering | $48.8 \pm 2.0^d$ | $50.0 \pm 1.0$ | 177 | N/A |
| Crystal<sup>a</sup> | 44.6 | $44.7 \pm 0.5$ | 142 | 1.44 |
| Jout-Jout1<sup>b</sup> | 44.6 | $44.6 \pm 0.5$ | 149 | 1.63 |
| Jout-Jout2<sup>b</sup> | 46.7 | $46.9 \pm 0.7$ | 176 | 1.30 |
| Jout-Jin<sup>b</sup> | 45.3 | $45.4 \pm 0.5$ | 151 | 1.37 |

<sup>a</sup> Calculated from the crystal structure
<sup>b</sup> Calculated from the final conformations of MD simulations
<sup>c</sup> Compared to experimental SAXS solution data
<sup>d</sup> Obtained from Guinier plot. The other $R_g$ values were obtained from GNOM.

**Table S3.** Thermal stability of the RIα_2_:J-PKAcα_2_ and RIα_2_:PKAcα_2_ holoenzymes

| Tm<sup>a</sup> (°C) | - MgATP | + MgATP |
| --- | --- | --- |
| RI$\alpha_2$:PKAc$\alpha_2$ | 52.93 ± 0.02 | 59.40 ± 0.03 |
| RI$\alpha_2$:J-PKAc$\alpha_2$ | 51.70 ± 0.01 | 58.90 ± 0.03 |
| ΔTm (wt/mutant) | 1.23 ± 0.02 | 0.50 ± 0.03 |

<sup>a</sup>Tm: temperature at which the protein denatures
All data are mean ± s.d. (n = 2 independent experiments).

**Figure S1.**
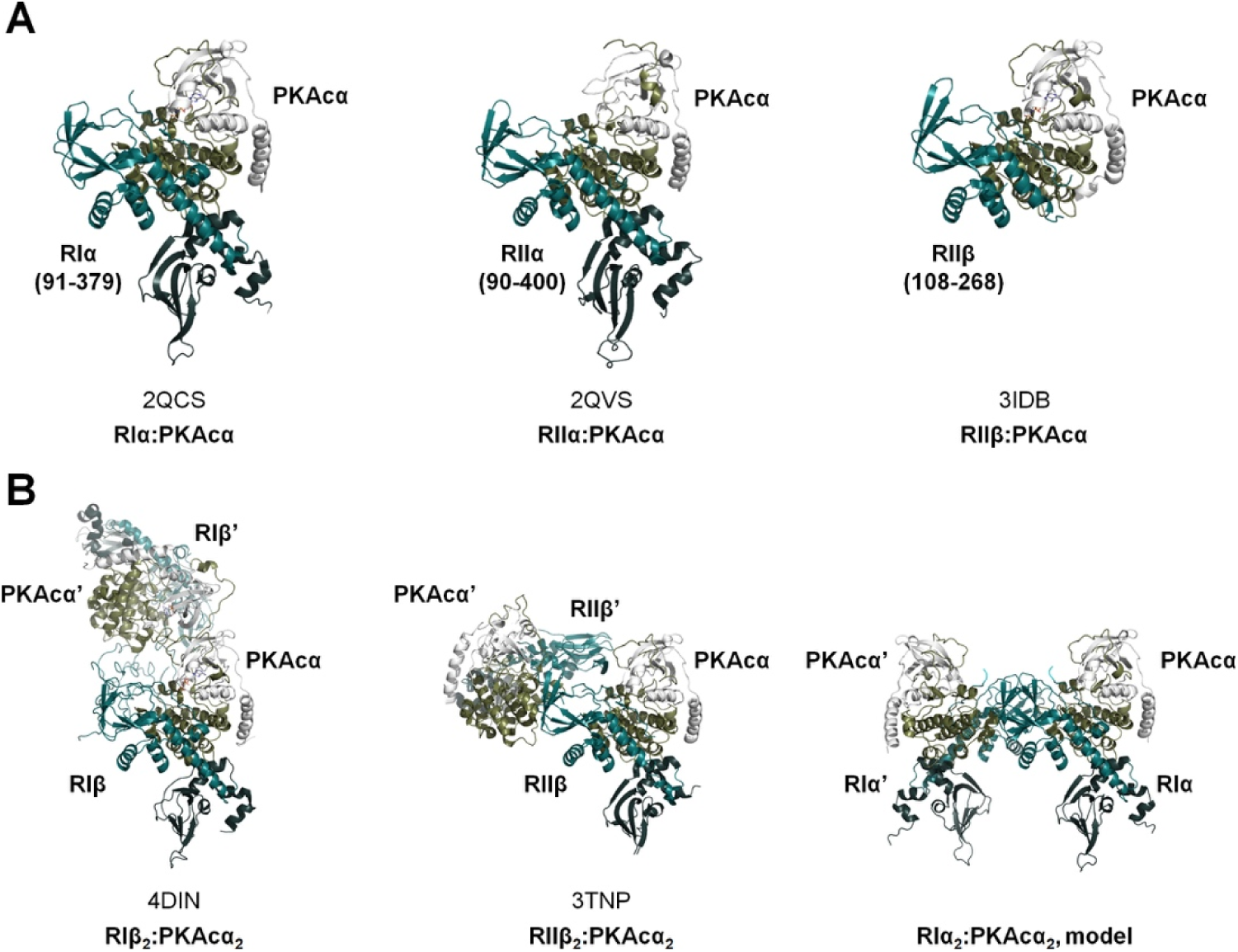
Summary of the previously determined structures of truncated PKA R:PKAcα heterodimers and wt R_2_:PKAcα_2_ holoenzymes. (A) Structures of truncated PKA R:PKAcα heterodimers: RIα (91-379):PKAcα (PDB ID 2QCS, left) (Kim et al., 2007), RIIα (90-400):PKAcα (PDB ID 2QVS, middle) (Wu et al., 2007) and RIIβ (108-268):PKAcα (PDB ID 3IDB, right) (Brown et al., 2009). (B) Structures of the wt R_2_:PKAcα_2_ holoenzymes: RIβ_2_:PKAcα_2_ (PDB ID 4DIN, left) (Ilouz et al., 2012), RIIβ_2_:PKAcα_2_ (PDB ID 3TNP, middle) (Zhang et al., 2012) holoenzymes which are determined by X-ray crystallography and the RIα_2_:PKAcα_2_ model (right) (Boettcher et al., 2011) based on crystal packing of two truncated RIα(73-244):PKAcα heterodimers. The right R:PKAcα heterodimers are aligned at the same orientation in all of the three holoenzymes.

**Figure S2.**
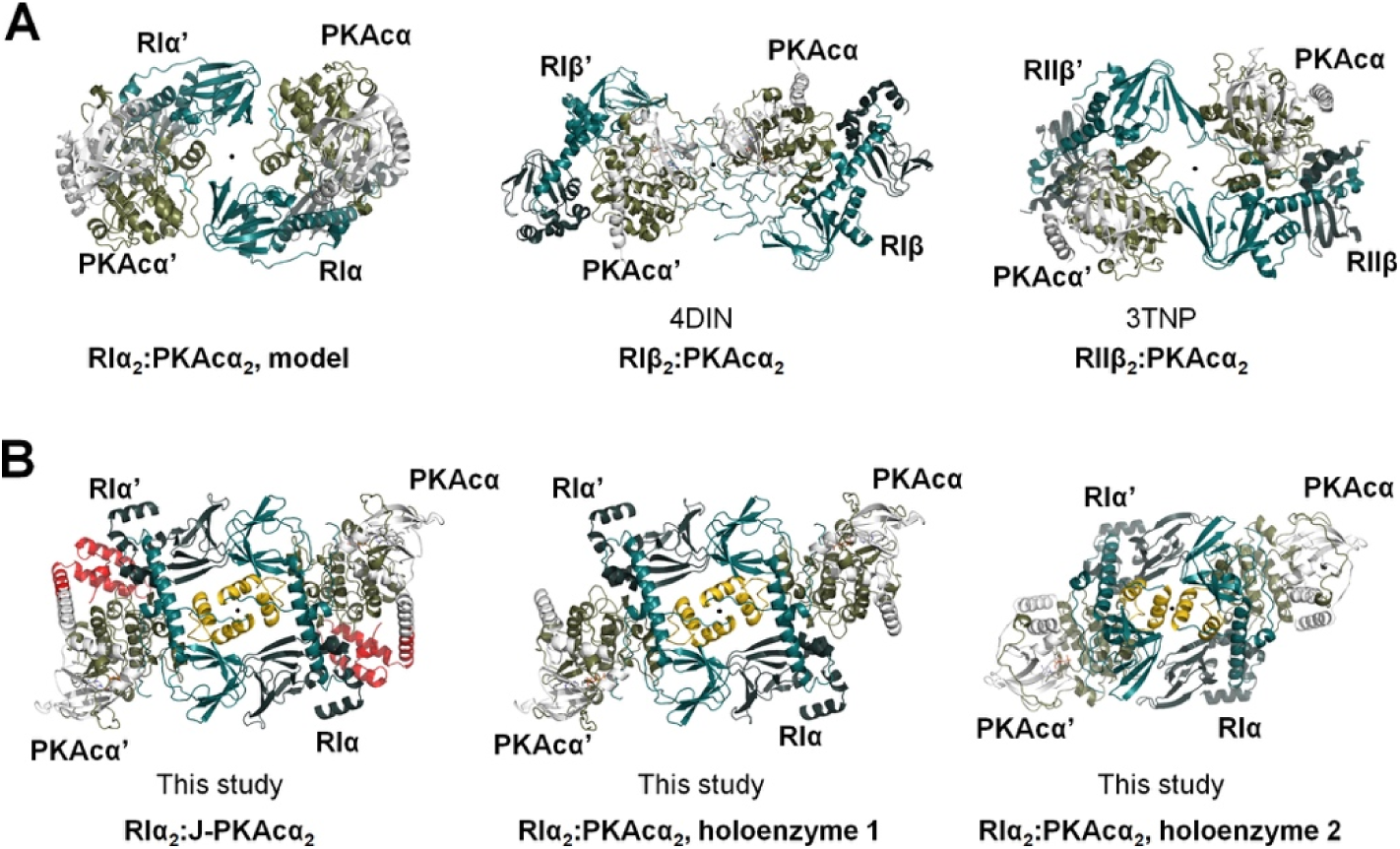
Bird-eye view of the structures of PKA R_2_:PKAcα_2_ and R_2_:J-PKAcα_2_ tetrameric holoenzymes determined previously and in this study. (A) Previously determined RIα_2_:PKAcα_2_ model (left) (Boettcher et al., 2011), RIβ_2_:PKAcα_2_ (PDB ID 4DIN, middle) (Ilouz et al., 2012), and RIIβ_2_:PKAcα_2_ (PDB ID 3TNP, right) (Zhang et al., 2012) tetrameric holoenzyme structures. (B) In this study determined RIα_2_:J-PKAcα_2_ and RIα_2_:J-PKAcα_2_ tetrameric holoenzyme structures. The twofold axis position for each holoenzyme is shown as a black dot in the middle..

**Figure S3.**
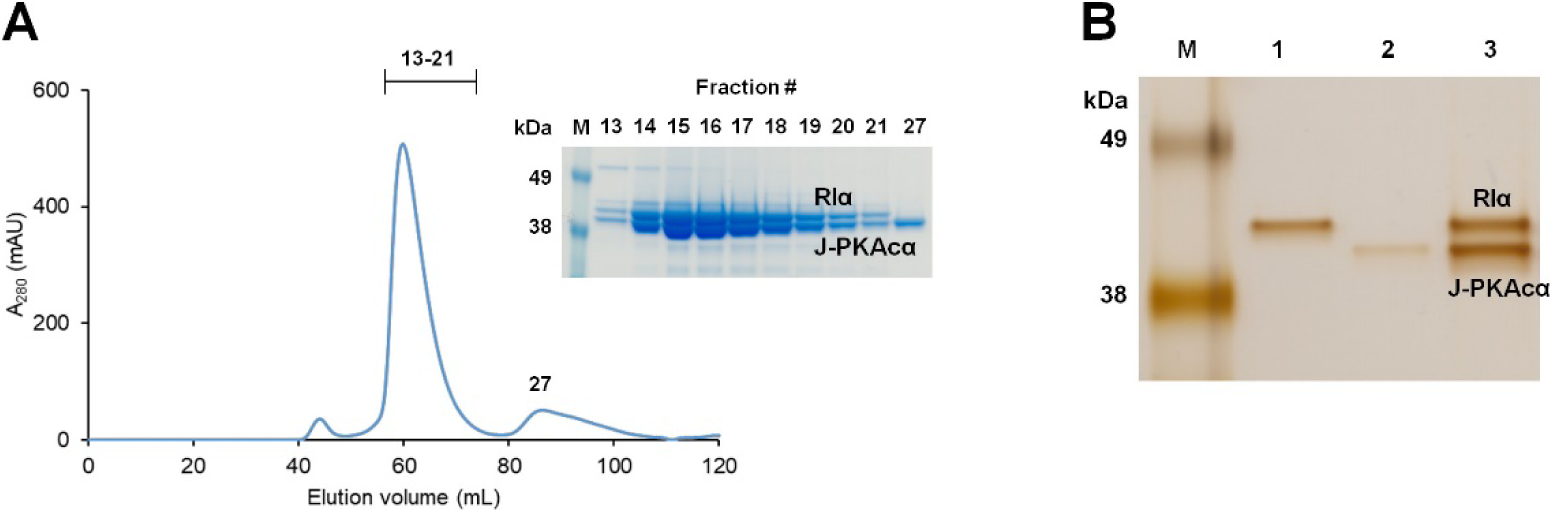
Formation of the chimeric RIα_2_:PKAcα_2_ holoenzyme. (A) Analytical gel filtration profile showing formation of RIα_2_:J-PKAcα_2_. (B) The diffracting crystals contain the full-length RIα_2_:J-PKAcα_2_ complex that was used for crystallization. The purified proteins RIα (lane 1), J-PKAcα (lane 2) and the dissolved diffracting crystals (lane 3) were run on a 7% tris-acetate SDS-PAGE gel and silver stained.

**Figure S4.**
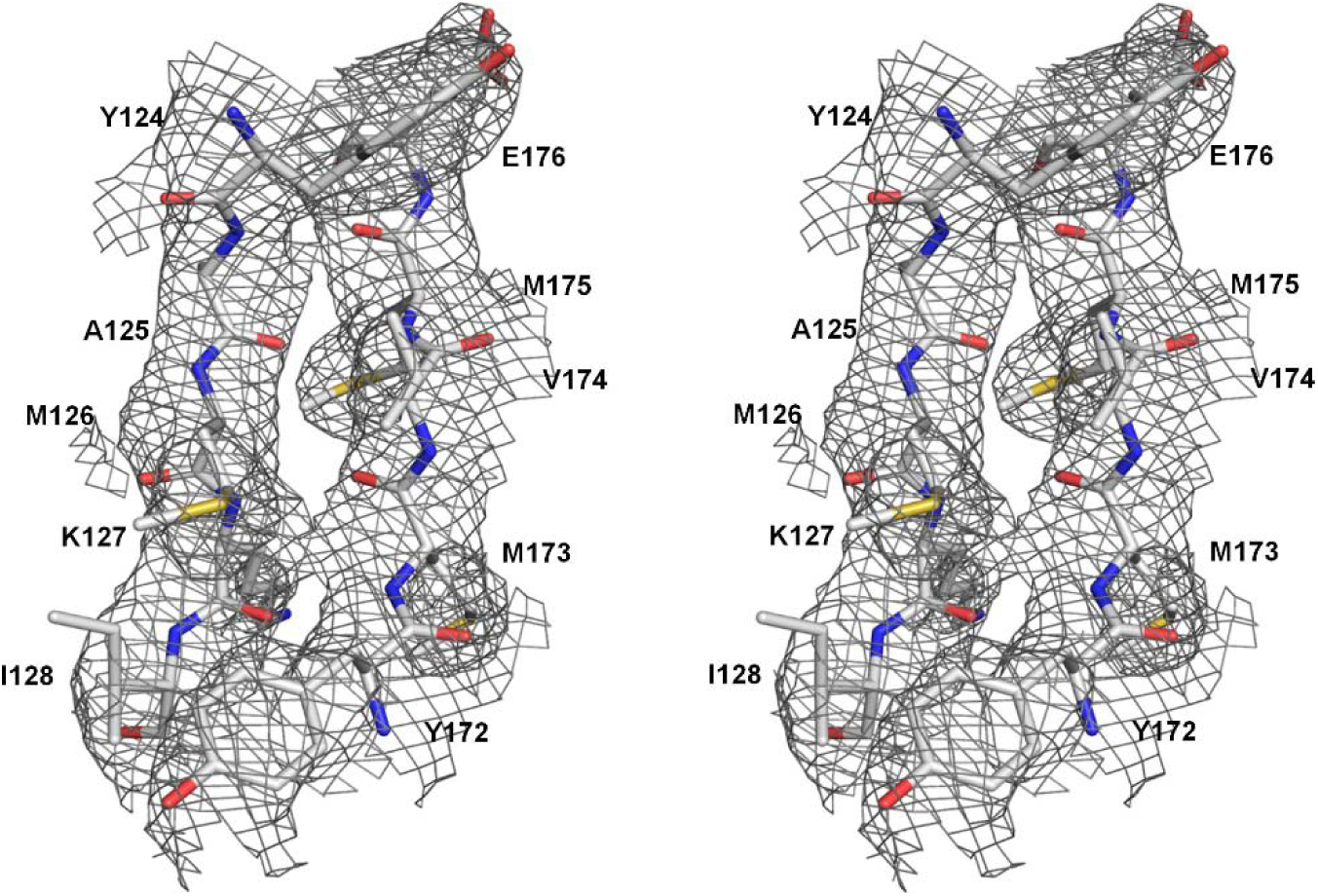
Cross-eyed stereo view of part of the RIα_2_:J-PKAcα_2_ holoenzyme structure in the 3.66 Å resolution *2Fo-Fc* map at 1 σ.

**Figure S5.**
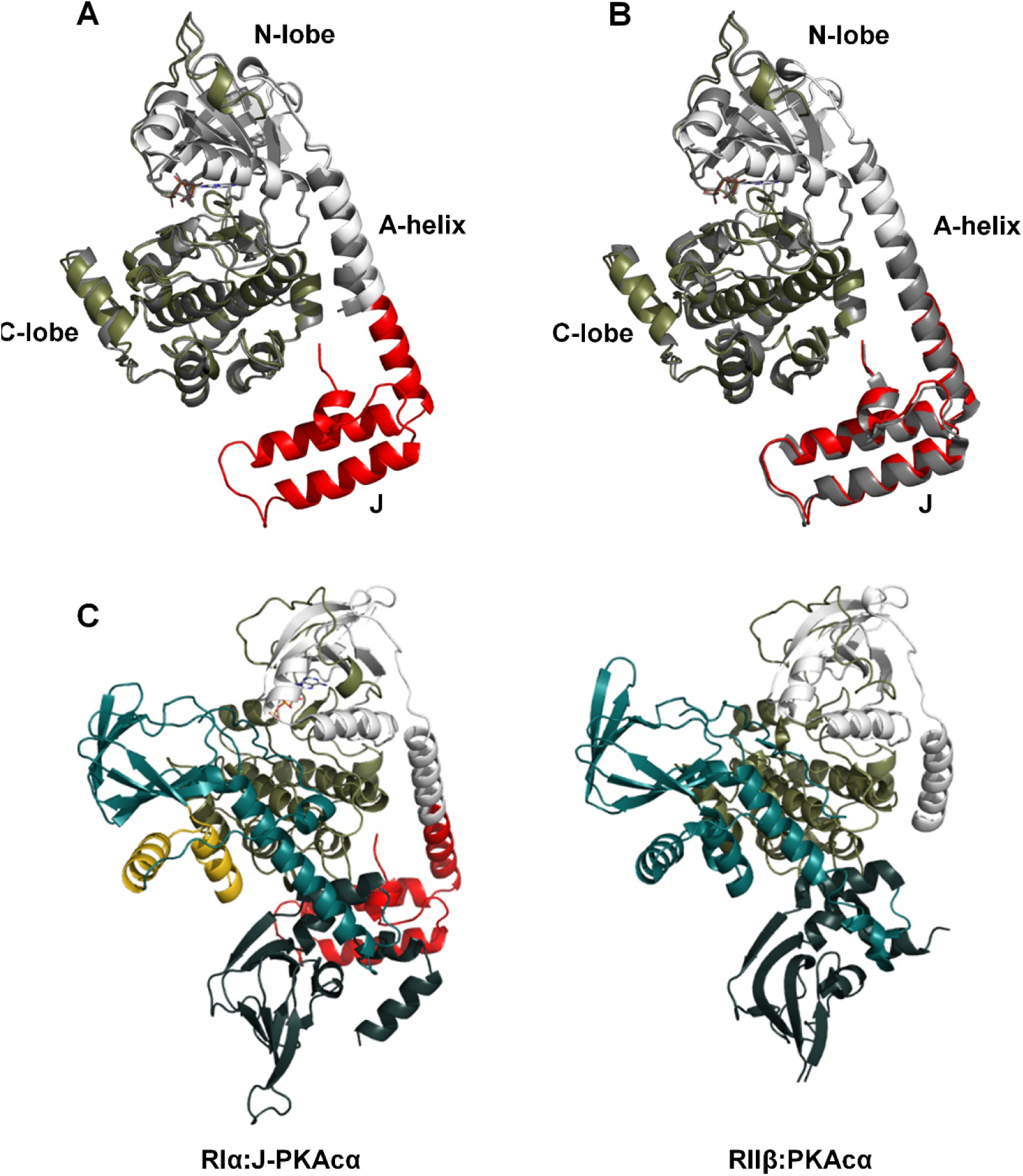
Addition of the J-domain does not significantly affect the structure of PKAcα and its binding with RIα. (A) Overlay of J-PKAcα (colored) in the chimeric holoenzyme and PKI-bound PKAcα (gray, PDB ID 1ATP). (B) Overlay of J-PKAcα in the chimeric holoenzyme (colored) and PKI-bound J-PKAcα (gray, PDB ID 4WB7). (C) Side-by-side comparison of RIα:J-PKAcα in the chimeric holoenzyme and a canonical R:PKAcα heterodimer in the RIIβ_2_:PKAcα_2_ holoenzyme (PDB ID 3TNP).

**Figure S6.**
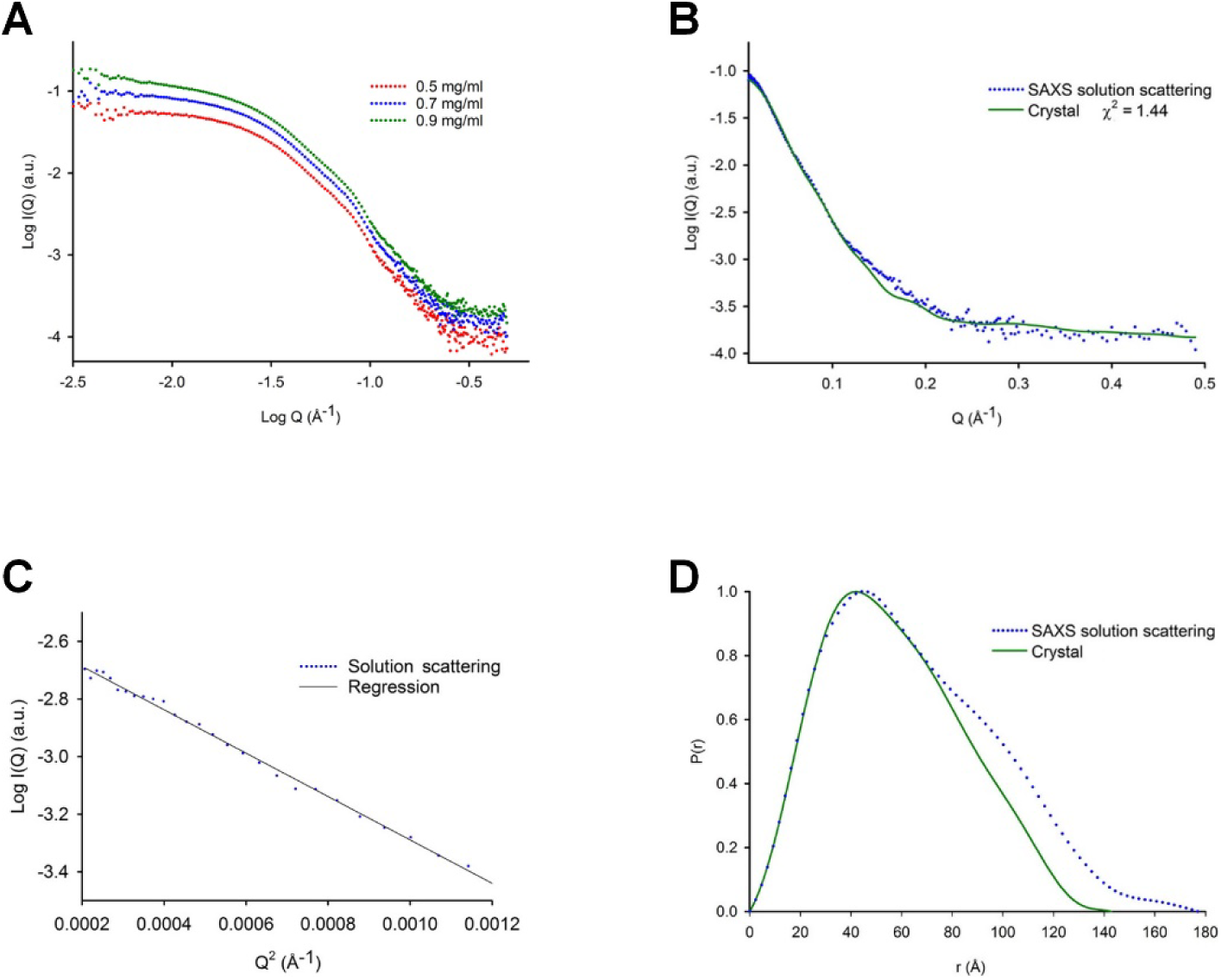
SAXS results from RIα_2_:J-PKAcα_2_. (A) SAXS profiles of RIα_2_:J-PKAcα_2_ at different concentrations. (B) Calculated scattering curve from crystal structure in continuous green line and SAXS experimental curve extrapolated to infinity dilution in blue dots. (C) Guinier plot, I_0_: 0.081, R_g_:48.8 ± 2.0 Å and Q_max_*R_g_: 1.26. (D) The P(r) functions from the crystal structure of RIα_2_:J-PKAcα_2_ in continuous solid green line and SAXS experimental data for the chimeric holoenzyme in blue dots.

**Figure S7.**
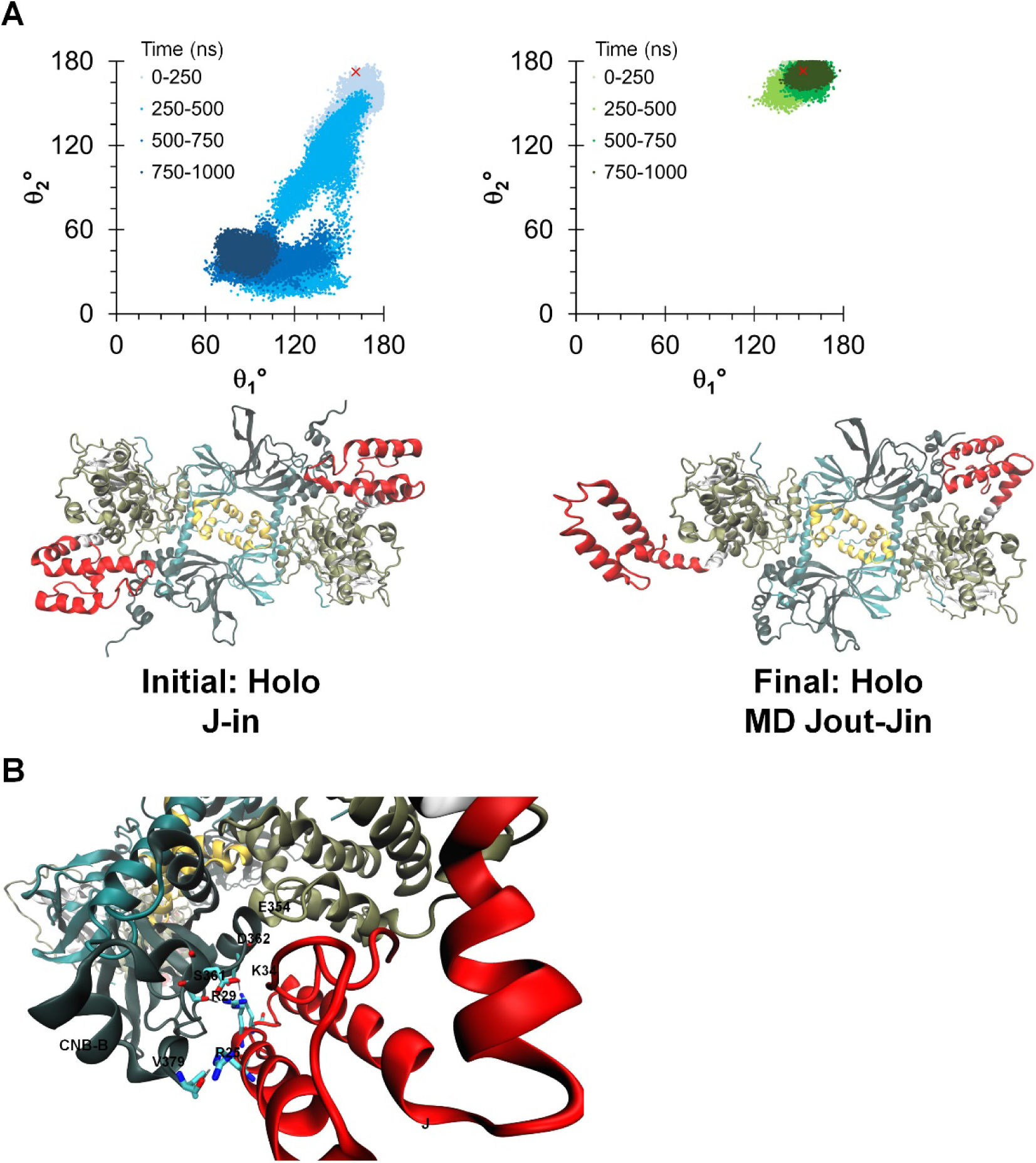
The J-domain configurations during the MD simulation started from the J-in state model. (A) Top: Simulation of RIα_2_:J-PKAcα_2_ showing the orientation of the J-domain for each copy of the chimera in the holoenzyme. The angles are the same as those defined in Figure 2. The red ‘x’ indicates the initial conformation of the J-domain. Bottom: Initial (left) and final (right) configurations of the J-domain in the holoenzyme. (B) Hydrogen bonds that formed between the J-domain and CNB-B domain during the simulation.

**Figure S8.**
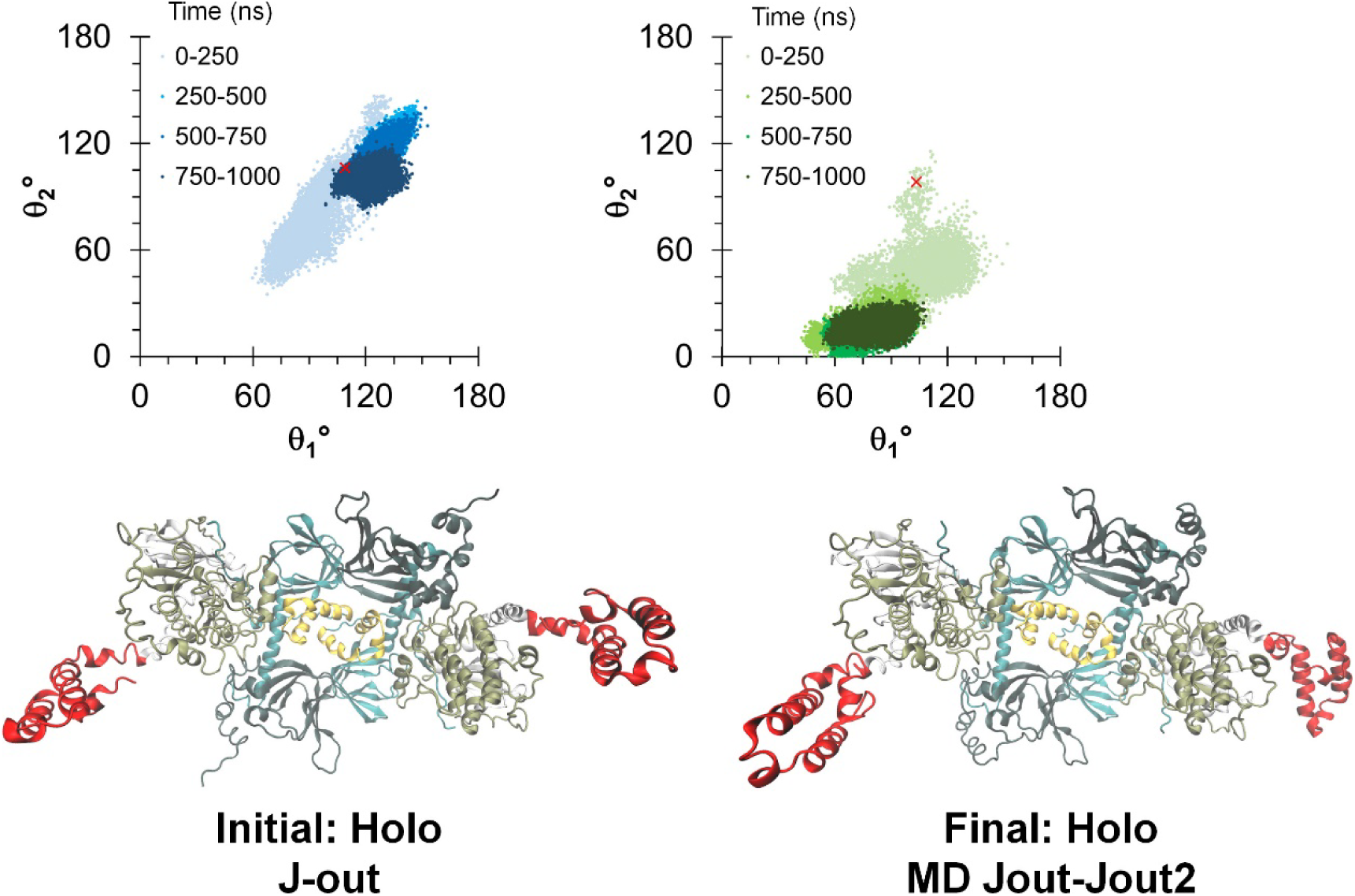
The J-domain configurations during the MD simulation started from the J-out state model. Top: Simulation of RIα_2_:J-PKAcα_2_ showing the orientation of the J-domain for each copy of the chimera in the holoenzyme. The angles are the same as those defined in Figure 2. The red ‘x’ indicates the initial conformation of the J-domain. Bottom: Initial (left) and final (right) configurations of the J-domain in the holoenzyme.

**Figure S9.**
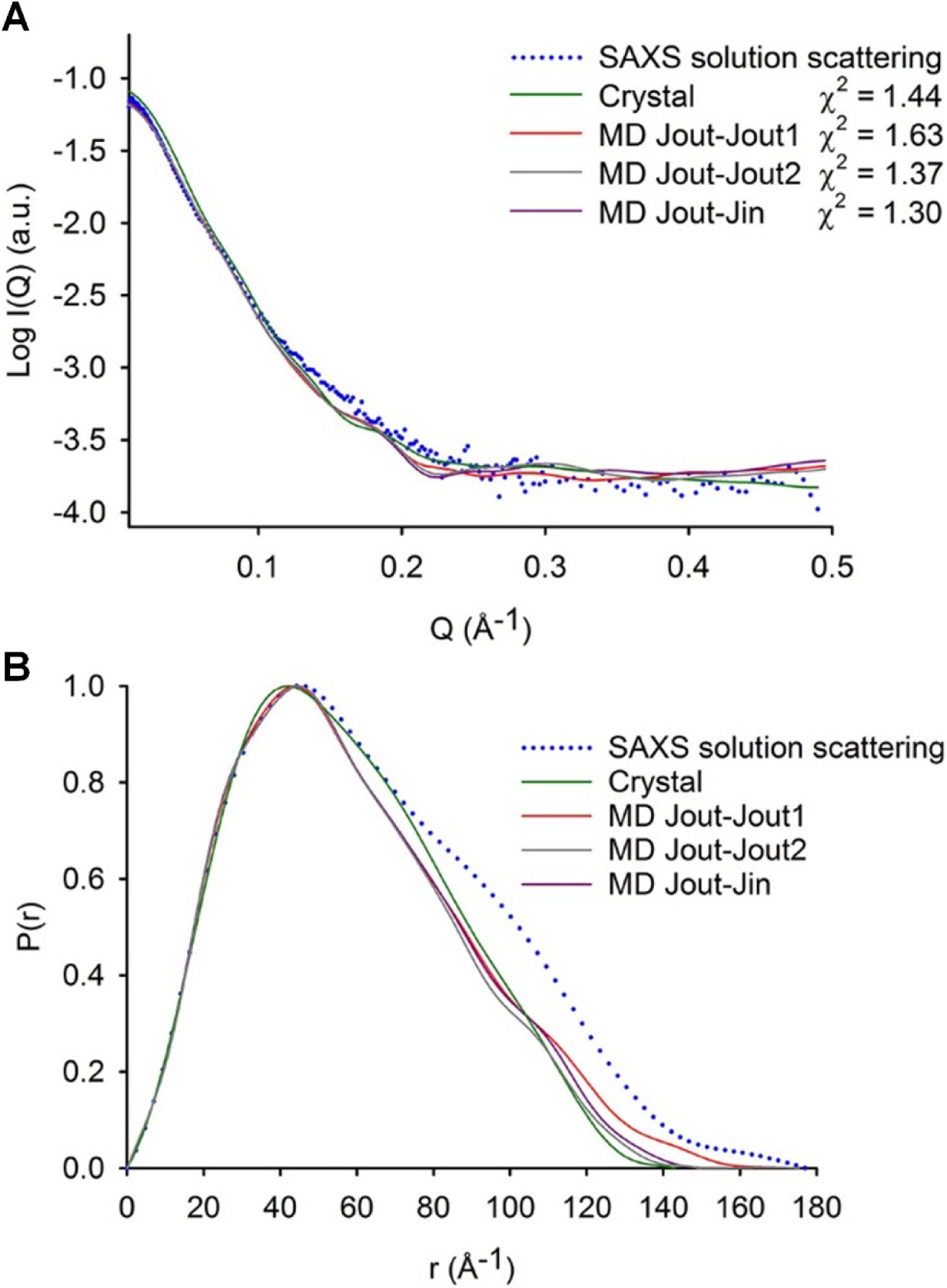
Comparison among the final conformations from holo MD simulations, holo crystal and SAXS experimental curves. (A) Calculated scattering curves from holo crystal and final conformations of MD simulations in solid lines and SAXS experimental curve extrapolated to infinity dilution in blue dots. (B) The P(r) functions from MD final conformations and holo crystal in solid lines and SAXS experimental data for the chimeric holoenzyme in blue dots. Calculated χ ^2^, R_g_ and D_max_ values are reported in Table S2.

**Figure S10.**
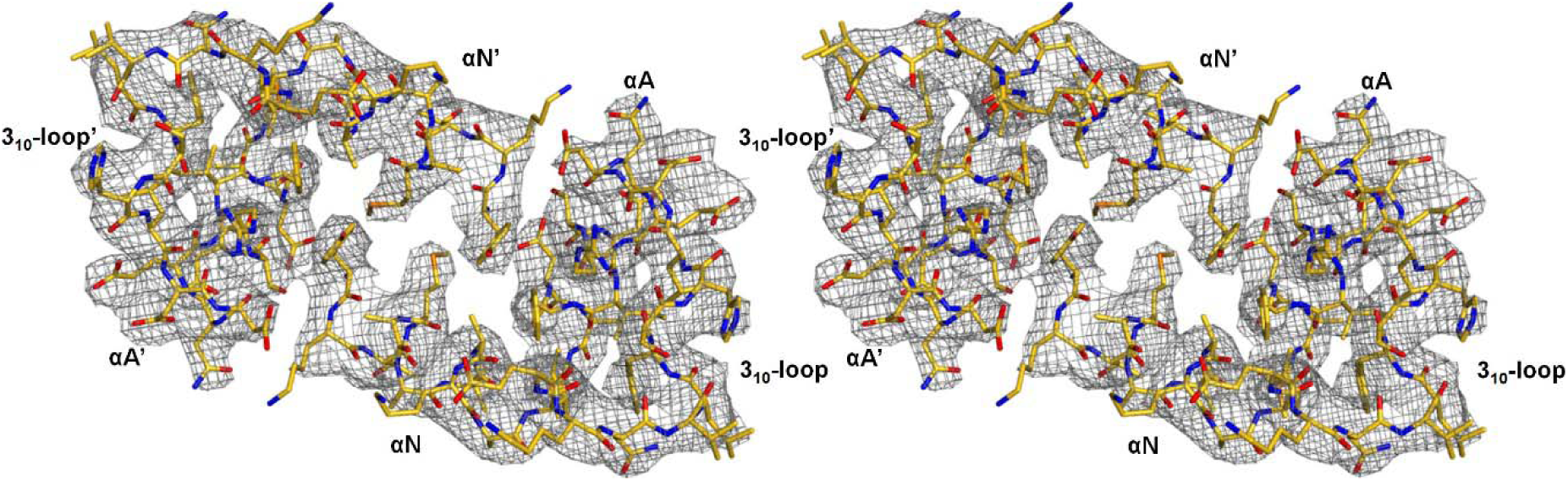
Cross-eyed stereo view of the N3A-N3A’ interface in the RIα_2_:J-PKAcα_2_ holoenzyme structure in the 3.66 Å resolution *2Fo-Fc* map at 1 σ.

**Figure S11.**
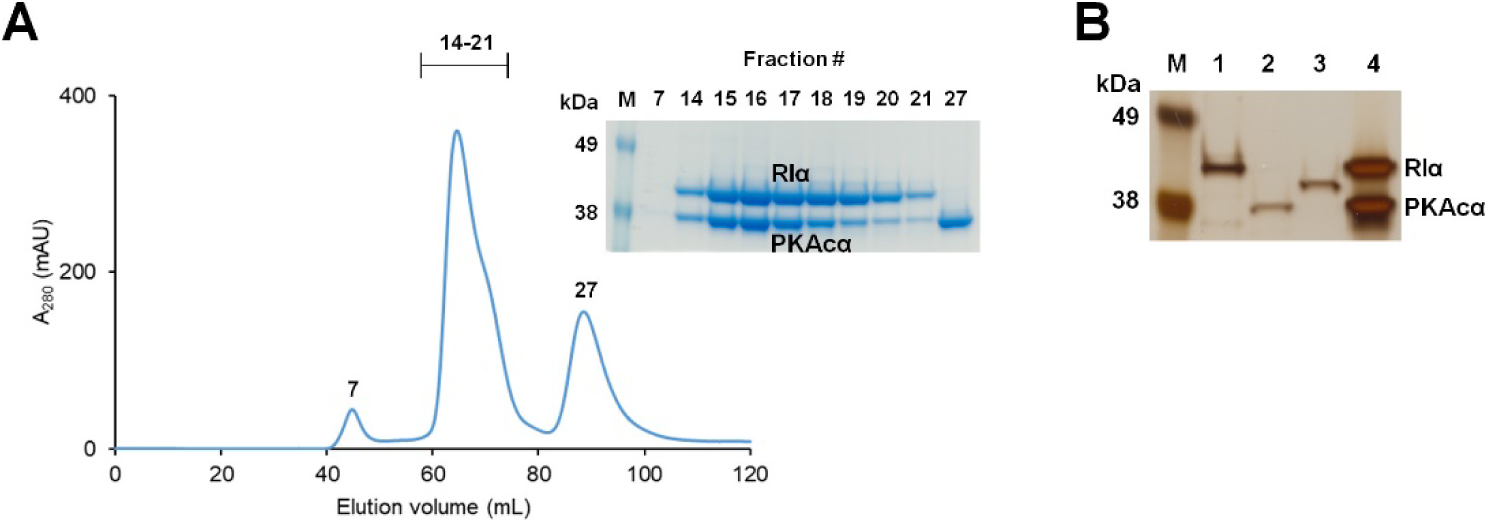
Formation of the wt RIα_2_:PKAcα_2_ holoenzyme. (A) Analytical gel filtration profile showing formation of RIα_2_:PKAcα_2_. (A) The diffracting crystals contain the full-length RIα_2_:PKAcα_2_ complex that was used for crystallization. The purified proteins RIα (lane 1), PKAcα (lane 2), J-PKAcα (lane 3) and the dissolved diffracting crystals (lane 4) were run on a 7% tris-acetate SDS-PAGE gel and silver stained.

**Figure S12.**
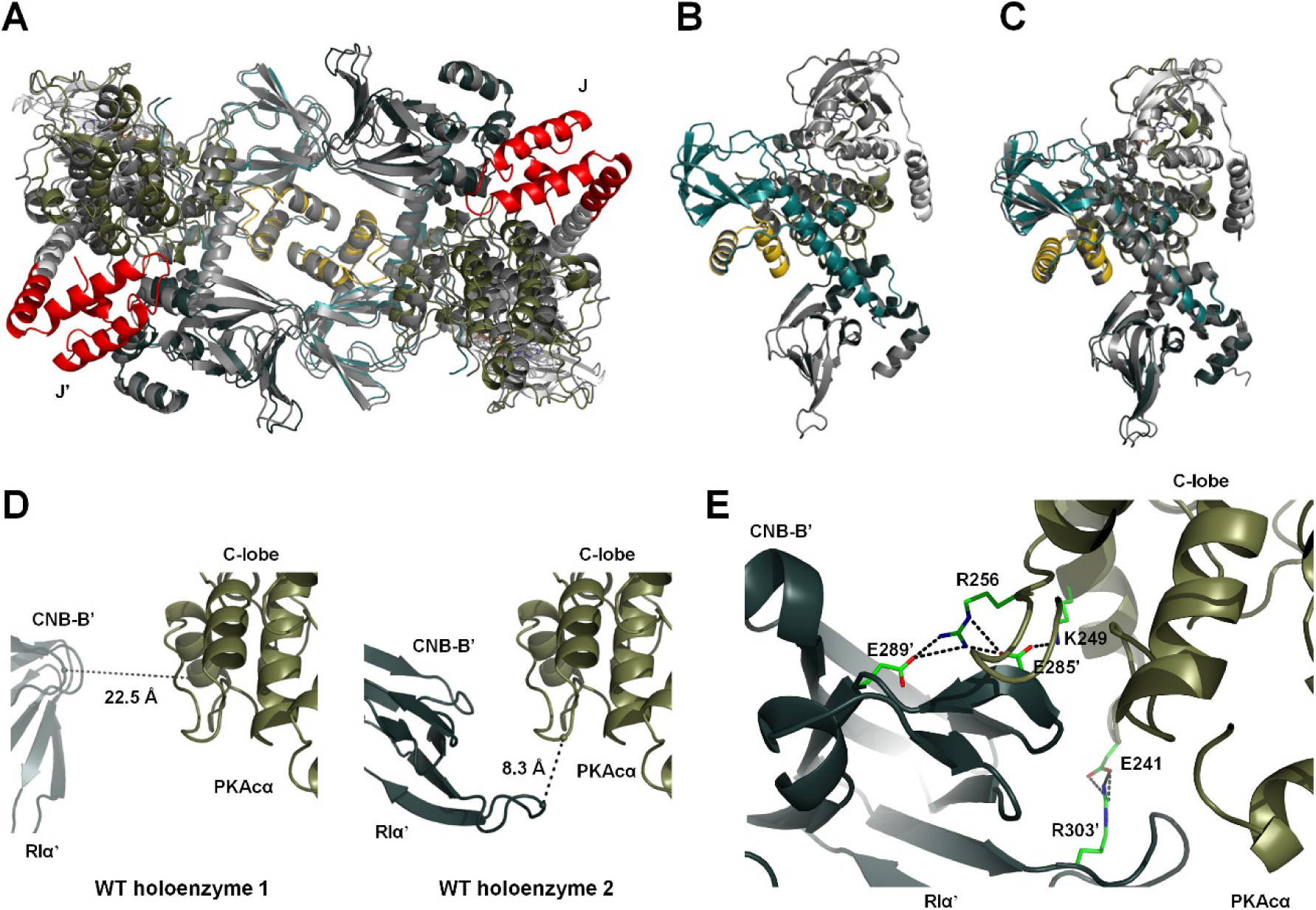
Wt holoenzyme 1 and 2. (A) Overlay of the RIα_2_:PKAcα_2_ holoenzyme 1 (gray) and chimeric RIα_2_:J-PKAcα_2_ holoenzyme (colored). (B) Overlay of the RIα:PKAcα heterodimers in the wt holoenzyme 1 (colored) and 2 (gray). (C) Overlay of RIα:PKAcα in the wt holoenzyme 1 (colored) and the previously reported RIα:PKAcα heterodimer (gray, PDB ID 2QCS). (D) The minimum Cα distances between PKAcα and RIα’ in wt holoenzyme 1 (left) and 2 (right). (E) Formation of salt bridges between the C-lobe of PKAcα and the CNB-B’ domain of RIα’ in wt holoenzyme 2 during MD simulation.

**Figure S13.**
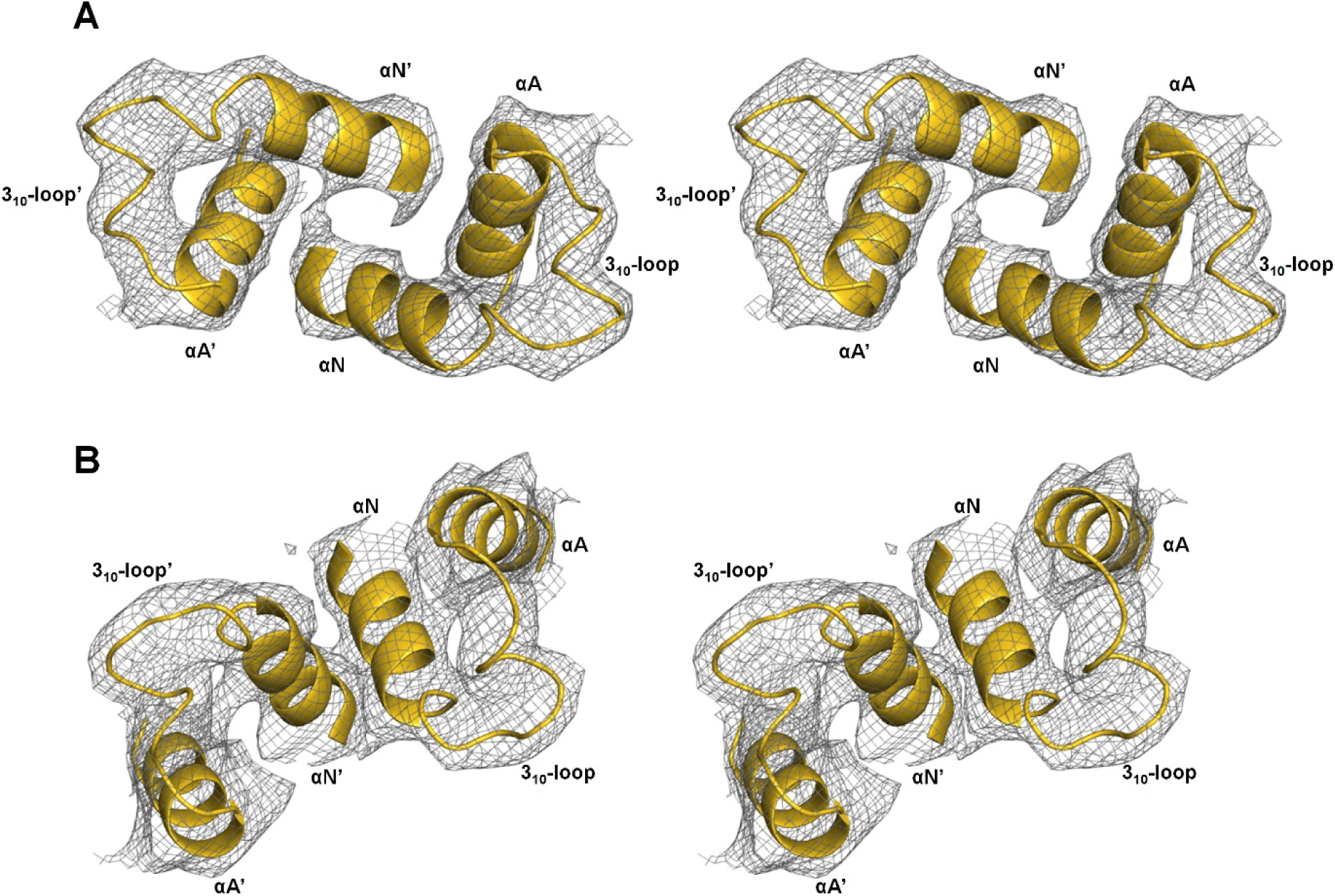
Cross-eyed stereo view of the N3A-N3A’ interfaces in the conformations 1 and 2 of the RIα_2_:J-PKAcα_2_ holoenzyme structure in the 3.66 Å resolution *2Fo-Fc* map at 1 σ, respectively. (A) Cross-eyed stereo view of the N3A-N3A’ interface in the conformation 1 of the RIα_2_:J-PKAcα_2_ holoenzyme structure. (B) Cross-eyed stereo view of the N3A-N3A’ interface in the conformation 2 of the RIα_2_:J-PKAcα_2_ holoenzyme structure.

**Figure S14.**
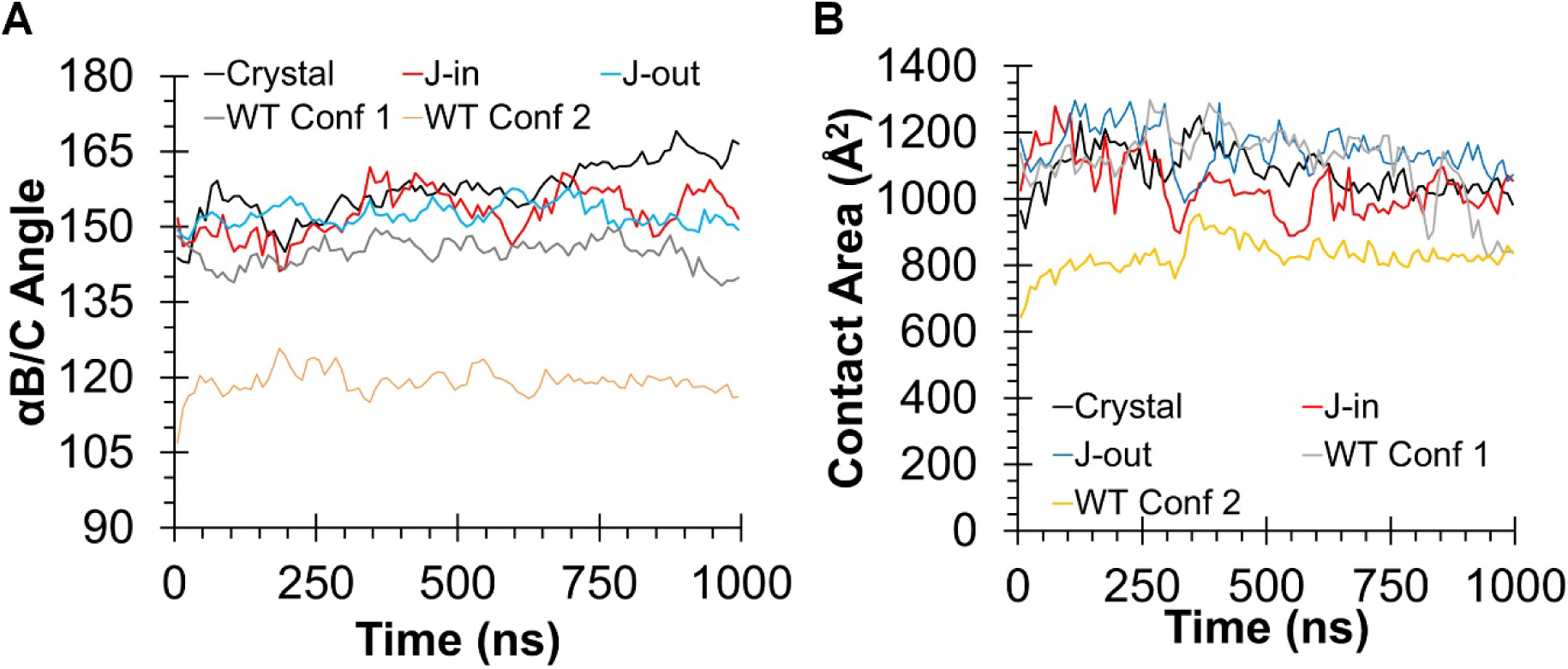
The RIα-RIα’ interfaces in the RIα chimeric holoenzyme (Crystal), models of the chimeric holoenzyme with both J-domains in J-in state (J-in) and with both J-domains in J-out state (J-out), and wt holo (WT) during MD simulations. (A) Orientation between the two αB/C-helices in the RIα dimers as a function of time. (B) Contact area of the RIα-RIα’ interfaces.

**Figure S15.**
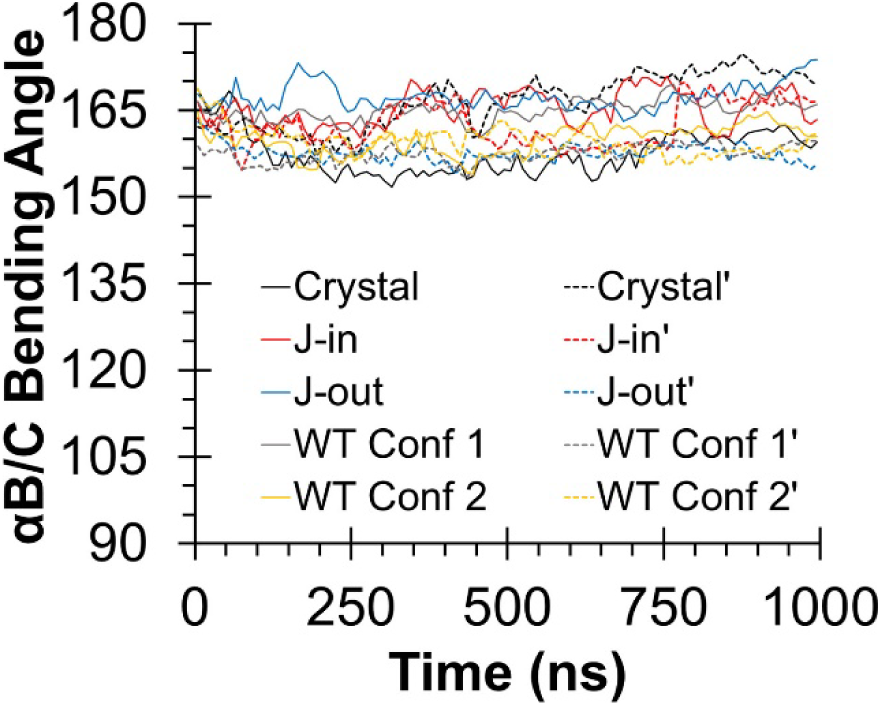
Dynamics of the αB/C-helix in the RIα chimeric holoenzyme (Crystal), models of the chimeric holoenzyme with both J-domains in J-in state (J-in) and with both J-domains in J-out state (J-out), and wt holo (WT) during MD simulations. Linearity of the αB/C-helices as defined by the Cα atoms of D225-G235-K250. The αB/C-helices in simulations do not sample the bent conformation that is observed in the cAMP-bound RIα homodimer. Solid lines indicate the αB/C bending angle in one R subunit while dashed lines indicate those in the symmetry-related R subunit in the holoenzyme.

## References

Adams, P. D., Afonine, P. V., Bunkóczi, G., Chen, V. B., Davis, I. W., Echols, N., Headd, J. J., Hung, L.W., Kapral, G. J., Grosse-Kunstleve, R. W., et al. (2010). PHENIX : a comprehensive Python-based system for macromolecular structure solution. Acta Crystallogr Sect D Biol Crystallogr. 66, 213–221.

Amieux, P.S., and McKnight, G.S. (2002). The essential role of RI alpha in the maintenance of regulated PKA activity. Ann. N. Y. Acad. Sci. 968, 75–95.

Amieux, P.S., Cummings, D.E., Motamed, K., Brandon, E.P., Wailes, L.A., Le, K., Idzerda, R.L., and Stanley McKnight, G. (1997). Compensatory regulation of RI?? protein levels in protein kinase A mutant mice. J. Biol. Chem. 272, 3993–3998.

Badireddy, S., Yunfeng, G., Ritchie, M., Akamine, P., Wu, J., Kim, C.W., Taylor, S.S., Qingsong, L., Swaminathan, K., and Anand, G.S. (2011). Cyclic AMP analog blocks kinase activation by stabilizing inactive conformation: conformational selection highlights a new concept in allosteric inhibitor design. Mol. Cell. Proteomics 10.

Boettcher, A.J., Wu, J., Kim, C., Yang, J., Bruystens, J., Cheung, N., Pennypacker, J.K., Blumenthal, D.A., Kornev, A.P., and Taylor, S.S. (2011). Realizing the Allosteric Potential of the Tetrameric Protein Kinase A RIα Holoenzyme. Structure 19, 265–276.

Brown, S.H.J., Wu, J., Kim, C., Alberto, K., and Taylor, S.S. (2009). Novel isoform-specific interfaces revealed by PKA RIIbeta holoenzyme structures. J. Mol. Biol. 393, 1070–1082.

Bruystens, J.G.H., Wu, J., Fortezzo, A., Kornev, A.P., Blumenthal, D.K., and Taylor, S.S. (2014). PKA RI homodimer structure reveals an intermolecular interface with implications for cooperative cAMP binding and Carney complex disease. Structure 22, 59–69.

Cheng, C.Y., Yang, J., Taylor, S.S., and Blumenthal, D.K. (2009). Sensing domain dynamics in protein kinase A-Iα complexes by solution x-ray scattering. J. Biol. Chem. 284, 35916–35925.

Cheung, J., Ginter, C., Cassidy, M., Franklin, M.C., Rudolph, M.J., Robine, N., Darnell, R.B., and Hendrickson, W.A. (2015). Structural insights into mis-regulation of protein kinase A in human tumors. Proc. Natl. Acad. Sci. U. S. A. 112, 1374–1379.

Craig, J.R., Peters, R.L., Edmondson, H.A., and Omata, M. (1980). Fibrolamellar carcinoma of the liver: A tumor of adolescents and young adults with distinctive clinico-pathologic features. Cancer 46, 372–379.

Darcy, D.G., Chiaroni-Clarke, R., Murphy, J.M., Honeyman, J.N., Bhanot, U., LaQuaglia, M.P., and Simon, S.M. (2015). The genomic landscape of fibrolamellar hepatocellular carcinoma: whole genome sequencing of ten patients. Oncotarget 6, 755–770.

Eggert, T., McGlynn, K.A., Duffy, A., Manns, M.P., Greten, T.F., and Altekruse, S.F. (2013). Epidemiology of fibrolamellar hepatocellular carcinoma in the USA, 2000-10. Gut 62, 1667–1668.

El-Serag, H.B., and Davila, J.A. (2004). Is fibrolamellar carcinoma different from hepatocellular carcinoma? A US population-based study. Hepatology 39, 798–803.

Engelholm, L.H., Riaz, A., Serra, D., Dagnæs-Hansen, F., Johansen, J. V., Santoni-Rugiu, E., Hansen, S.H., Niola, F., and Frödin, M. (2017). CRISPR/Cas9 Engineering of Adult Mouse Liver Demonstrates That the Dnajb1 – Prkaca Gene Fusion Is Sufficient to Induce Tumors Resembling Fibrolamellar Hepatocellular Carcinoma. Gastroenterology 153, 1662–1673.

Graham, R., Lackner, K., Terracciano, L., González-Cantú, Y., Maleszewski, J.J., Greipp, P.T., Simon, S.M., and Torbenson, M.S. (2017). Fibrolamellar Carcinoma in the Carney Complex: PRKAR1A Loss Instead of the Classic DNAJB1-PRKACA Fusion. Hepatology.

Greene, E.L., Horvath, A.D., Nesterova, M., Giatzakis, C., Bossis, I., and Stratakis, C.A. (2008). In vitro functional studies of naturally occurring pathogenic PRKAR1A mutations that are not subject to nonsense mRNA decay. Hum. Mutat. 29, 633–639.

Hirakis, S.P., Malmstrom, R.D., and Amaro, R.E. (2017). Molecular Simulations Reveal an Unresolved Conformation of the Type IA Protein Kinase A Regulatory Subunit and Suggest Its Role in the cAMP Regulatory Mechanism. Biochemistry 56, 3885–3888.

Honeyman, J.N., Simon, E.P., Robine, N., Chiaroni-Clarke, R., Darcy, D.G., Lim, I.I.P., Gleason, C.E., Murphy, J.M., Rosenberg, B.R., Teegan, L., et al. (2014). Detection of a Recurrent DNAJB1-PRKACA Chimeric Transcript in Fibrolamellar Hepatocellular Carcinoma. Science (80-.). 343, 1010–1014.

Ilouz, R., Bubis, J., Wu, J., Yim, Y.Y., Deal, M.S., Kornev, A.P., Ma, Y., Blumenthal, D.K., and Taylor, S.S. (2012). Localization and quaternary structure of the PKA RIβ holoenzyme. Proc. Natl. Acad. Sci. U. S. A. 109, 12443–12448.

Kakar, S., Burgart, L.J., Batts, K.P., Garcia, J., Jain, D., and Ferrell, L.D. (2005). Clinicopathologic features and survival in fibrolamellar carcinoma: Comparison with conventional hepatocellular carcinoma with and without cirrhosis. Mod. Pathol. 18, 1417–1423.

Kastenhuber, E.R., Lalazar, G., Houlihan, S.L., Tschaharganeh, D.F., Baslan, T., Chen, C.-C., Requena, D., Tian, S., Bosbach, B., Wilkinson, J.E., et al. (2017). DNAJB1-PRKACA fusion kinase interacts with β-catenin and the liver regenerative response to drive fibrolamellar hepatocellular carcinoma. Proc. Natl. Acad. Sci. U. S. A. 13076–13084.

Katzenstein, H.M., Krailo, M.D., Malogolowkin, M.H., Ortega, J.A., Qu, W., Douglass, E.C., Feusner, J.H., Reynolds, M., Quinn, J.J., Newman, K., et al. (2003). Fibrolamellar hepatocellular carcinoma in children and adolescents. Cancer 97, 2006–2012.

Kim, C., Cheng, C.Y., Saldanha, S.A., and Taylor, S.S. (2007). PKA-I holoenzyme structure reveals a mechanism for cAMP-dependent activation. Cell 130, 1032–1043.

Lalazar, G., and Simon, S.M. (2018). Fibrolamellar Carcinoma: Recent Advances and Unresolved Questions on the Molecular Mechanisms. Semin. Liver Dis. 38, 51–59.

Li, F., Milind Gangal, Jones, J.M., Jason Deich, Kimberly E. Lovett, Susan S. Taylor, A., and David A. Johnson (2000). Consequences of cAMP and Catalytic-Subunit Binding on the Flexibility of the A-Kinase Regulatory Subunit. 39, 15626–15632.

Lim, I., Farber, B., and LaQuaglia, M. (2014). Advances in Fibrolamellar Hepatocellular Carcinoma: A Review. Eur. J. Pediatr. Surg. 24, 461–466.

Linglart, A., Fryssira, H., Hiort, O., Holterhus, P.-M., Perez de Nanclares, G., Argente, J., Heinrichs, C., Kuechler, A., Mantovani, G., Leheup, B., et al. (2012). *PRKAR1A* and *PDE4D* Mutations Cause Acrodysostosis but Two Distinct Syndromes with or without GPCR-Signaling Hormone Resistance. J. Clin. Endocrinol. Metab. 97, E2328–E2338.

Mavros, M.N., Mayo, S.C., Hyder, O., and Pawlik, T.M. (2012). A Systematic Review: Treatment and Prognosis of Patients with Fibrolamellar Hepatocellular Carcinoma. J. Am. Coll. Surg. 215, 820–830.

Oikawa, T., Wauthier, E., Dinh, T.A., Selitsky, S.R., Reyna-Neyra, A., Carpino, G., Levine, R., Cardinale, V., Klimstra, D., Gaudio, E., et al. (2015). Model of fibrolamellar hepatocellular carcinomas reveals striking enrichment in cancer stem cells. Nat. Commun. 6, 8070.

P. Barros, E., Malmstrom, R.D., Nourbakhsh, K., Del Rio, J.C., Kornev, A.P., Taylor, S.S., and Amaro, R.E. (2017). Electrostatic Interactions as Mediators in the Allosteric Activation of Protein Kinase A RIα. Biochemistry 56, 1536–1545.

Park, K.U., Kim, H.-S., Lee, S.K., Jung, W.-W., and Park, Y.-K. (2012). Novel Mutation in PRKAR1A in Carney Complex. Korean J. Pathol. 46, 595–600.

Riggle, K.M., Riehle, K.J., Kenerson, H.L., Turnham, R., Homma, M.K., Kazami, M., Samelson, B., Bauer, R., McKnight, G.S., Scott, J.D., et al. (2016a). Enhanced cAMP-stimulated protein kinase A activity in human fibrolamellar hepatocellular carcinoma. Pediatr. Res. 80, 110–118.

Riggle, K.M., Turnham, R., Scott, J.D., Yeung, R.S., and Riehle, K.J. (2016b). Fibrolamellar Hepatocellular Carcinoma: Mechanistic Distinction From Adult Hepatocellular Carcinoma. Pediatr. Blood Cancer 63, 1163–1167.

Simon, E.P., Freije, C.A., Farber, B.A., Lalazar, G., Darcy, D.G., Honeyman, J.N., Chiaroni-Clarke, R., Dill, B.D., Molina, H., Bhanot, U.K., et al. (2015). Transcriptomic characterization of fibrolamellar hepatocellular carcinoma. Proc. Natl. Acad. Sci. U. S. A. 112, E5916–E5925.

Søberg, K., Moen, L.V., Skålhegg, B.S., and Laerdahl, J.K. (2017). Evolution of the cAMP-dependent protein kinase (PKA) catalytic subunit isoforms. PLoS One.

Taylor, S.S., Ilouz, R., Zhang, P., and Kornev, A.P. (2012). Assembly of allosteric macromolecular switches: lessons from PKA. Nat. Rev. Mol. Cell Biol. 13, 646–658.

Terracciano, L.M., Tornillo, L., Avoledo, P., Von Schweinitz, D., Kühne, T., and Bruder, E. (2004). Fibrolamellar hepatocellular carcinoma occurring 5 years after hepatocellular adenoma in a 14-year-old girl: a case report with comparative genomic hybridization analysis. Arch. Pathol. Lab. Med. 128, 222–226.

Tomasini, M.D., Wang, Y., Karamafrooz, A., Li, G., Beuming, T., Gao, J., Taylor, S.S., Veglia, G., and Simon, S.M. (2018). Conformational Landscape of the PRKACA-DNAJB1 Chimeric Kinase, the Driver for Fibrolamellar Hepatocellular Carcinoma. Sci. Rep. 8, 720.

Torbenson, M. (2012). Fibrolamellar carcinoma: 2012) update. Scientifica (Cairo). 2012, 743790.

Veugelers, M., Wilkes, D., Burton, K., McDermott, D.A., Song, Y., Goldstein, M.M., La Perle, K., Vaughan, C.J., O’Hagan, A., Bennett, K.R., et al. (2004). Comparative PRKAR1A genotypephenotype analyses in humans with Carney complex and prkar1a haploinsufficient mice. Proc. Natl. Acad. Sci. U. S. A. 101, 14222–14227.

Weeda, V.B., Murawski, M., McCabe, A.J., Maibach, R., Brugières, L., Roebuck, D., Fabre, M., Zimmermann, A., Otte, J.B., Sullivan, M., et al. (2013). Fibrolamellar variant of hepatocellular carcinoma does not have a better survival than conventional hepatocellular carcinoma--results and treatment recommendations from the Childhood Liver Tumour Strategy Group (SIOPEL) experience. Eur. J. Cancer 49, 2698–2704.

Wu, J., Jones, J.M., Xuong, N.H., Ten Eyck, L.F., and Taylor, S.S. (2004a). Crystal structures of RIα subunit of cyclic adenosine 5′-monophosphate (cAMP)-dependent protein kinase complexed with (R p)-adenosine 3′, 5′-cyclic monophosphothioate, the phosphothioate analogues of cAMP. Biochemistry 43, 6620–6629.

Wu, J., Brown, S., Xuong, N.-H., and Taylor, S.S. (2004b). RIalpha subunit of PKA: a cAMP-free structure reveals a hydrophobic capping mechanism for docking cAMP into site B. Structure 12, 1057–1065.

Wu, J., Brown, S.H.J., von Daake, S., and Taylor, S.S. (2007). PKA type IIalpha holoenzyme reveals a combinatorial strategy for isoform diversity. Science 318, 274–279.

Zhang, P., Smith-Nguyen, E. V., Keshwani, M.M., Deal, M.S., Kornev, A.P., and Taylor, S.S. (2012). Structure and Allostery of the PKA RII Tetrameric Holoenzyme. Science (80). 335, 712–716.

Zhang, P., Ye, F., Bastidas, A.C., Kornev, A.P., Wu, J., Ginsberg, M.H., and Taylor, S.S. (2015). An Isoform-Specific Myristylation Switch Targets Type II PKA Holoenzymes to Membranes. Structure.

Zheng, J., Trafny, E.A., Knighton, D.R., Xuong, N.H., Taylor, S.S., Ten Eyck, L.F., and Sowadski, J.M. (1993). 2.2 A refined crystal structure of the catalytic subunit of cAMP-dependent protein kinase complexed with MnATP and a peptide inhibitor. Acta Crystallogr. D. Biol. Crystallogr. 49, 362–365.

